# CITEseq analysis of non-small-cell lung cancer lesions reveals an axis of immune cell activation associated with tumor antigen load and *TP53* mutations

**DOI:** 10.1101/2020.07.16.207605

**Authors:** Andrew M. Leader, John A. Grout, Christie Chang, Barbara Maier, Alexandra Tabachnikova, Laura Walker, Alona Lansky, Jessica LeBerichel, Naussica Malissen, Melanie Davila, Jerome Martin, Giuliana Magri, Kevin Tuballes, Zhen Zhao, Francesca Petralia, Robert Samstein, Natalie Roy D’Amore, Gavin Thurston, Alice Kamphorst, Andrea Wolf, Raja Flores, Pei Wang, Mary Beth Beasley, Helene Salmon, Adeeb H. Rahman, Thomas U. Marron, Ephraim Kenigsberg, Miriam Merad

**Author notes:** Authors contributed equally to this work (Ephraim Kenigsberg, Miriam Merad). Program for Inflammatory and Cardiovascular Disorders, Institut Hospital del Mar d’Investigacions Mèdiques (IMIM), 08003 Barcelona, Spain. INSERM U932, Institut Curie, 26 rue d’Ulm, 75005 Paris, France (Helene Salmon).

## Abstract

Immunotherapy is becoming a mainstay in the treatment of NSCLC. While tumor mutational burden (TMB) has been shown to correlate with response to immunotherapy, little is known about the relation of the baseline immune response with the tumor genotype. Here, we profiled 35 early stage NSCLC lesions using multiscale single cell sequencing. Unsupervised clustering identified in a subset of patients a key cellular module consisting of *PDCD1+ CXCL13*+ activated T cells, IgG+ plasma cells, and *SPP1*+ macrophages, referred to as the lung cancer activation module (LCAM^hi^). Transcriptional data from two NSCLC cohorts confirmed a subset of patients with LCAM^hi^ enrichment, which was independent of overall immune cell content. The LCAM^hi^ module strongly correlated with TMB, expression of cancer testis antigens, and with *TP53* mutations in smokers and non-smokers. These data establish LCAM as a key mode of immune cell activation associated with high tumor antigen load and driver mutations.

## INTRODUCTION

Lung cancer is the most common cause of cancer-related death^1^, and the most common subgroup of lung cancer is non-small cell lung cancer (NSCLC)^2^. In recent years, immune checkpoint blockade (ICB) targeting the PD-1/PD-L1 axis has become first-line therapy for a majority of patients with metastatic and locally advanced disease^3^. Though ICB studies have achieved improved overall survival, fewer than half of patients achieve significant clinical benefit, though still may experience physical and financial toxicity. Biomarkers are lacking to determine optimal treatment regimens for patients, as our understanding of tumor-associated immune phenotypes and immune correlates of response to ICB remains incomplete.

While multiple studies have used single-cell assays to profile NSCLC tumor-infiltrating immune cells in comparison to patient-matched, non-involved lung (nLung)^4,5^, blood^6^, or both^7,8^, or characterized tumor-infiltrating lymphocytes (TIL)^9-12^, we continue to lack a comprehensive understanding of how immune cell phenotypes vary across patients. In particular, it remains unclear which immune cell populations and phenotypes are associated with robust, tumor-directed T cell responses and response to ICB, and how these features are connected to tumor-cell intrinsic characteristics such as tumor mutational burden (TMB)^13,14^. A deeper analysis is further required for uncovering the cell types and states associated with immunostimulatory versus immunoregulatory presentation of tumor-associated antigens, as well as parsing the tumor-related effects on tissue-resident and migratory innate cell types. Attempts to integrate these data across the innate and adaptive arms of the immune system are of crucial importance to optimizing rational design of immunotherapies. Furthermore, while response to ICB has been associated with specific patient groups, individual driver mutations, the degree of immune infiltrate, and TMB, the complex interplay between these factors remains poorly understood.

Here, we sought to define the molecular immune states induced in the tumor microenvironment by profiling an expanded patient cohort compared to previous related studies^5,6^ via multiscale single-cell analyses. We integrated the results of single-cell RNA sequencing (scRNAseq) of immune cells with cellular indexing of transcriptomes and epitopes by sequencing (CITEseq)^15^, a method allowing for combined scRNAseq and multiplexed single-cell surface protein measurement. To further elucidate the TCR landscape across T cell phenotypes, we analyzed these results together with joint scRNAseq/TCRseq. We revealed a pattern of inter-tumor variability involving innate and adaptive immune responses which we validated in two bulk RNA datasets, allowing us to detect an association with TMB and tumor driver mutations.

## RESULTS

### Integrative analyses unify phenotypic mappings across substrates and datasets

To probe transcriptional states of immune cells in the lung cancer microenvironment, we set out to profile cells from a cohort of untreated, early-stage NSCLC patients undergoing resection with curative intent (Figure 1A). The cohort was diverse with respect to age, smoking status, sex, and histological subtype (Figure 1B). We generated three datasets integrating antibody profiling of surface marker proteins using CITEseq^15^, scRNAseq, and TCRseq with single cell resolution (Tables S1-S3). We performed CITEseq on matched tumor and non-involved lung (nLung) tissues from 7 patients, in addition to performing scRNAseq of matched tumor and nLung in 28 additional patients. Finally, to expand on our annotation of T cell clusters based on the distribution of clonally expanded populations, we performed paired single-cell TCRseq and scRNAseq on T cells isolated from 3 patients.

**Figure 1.**
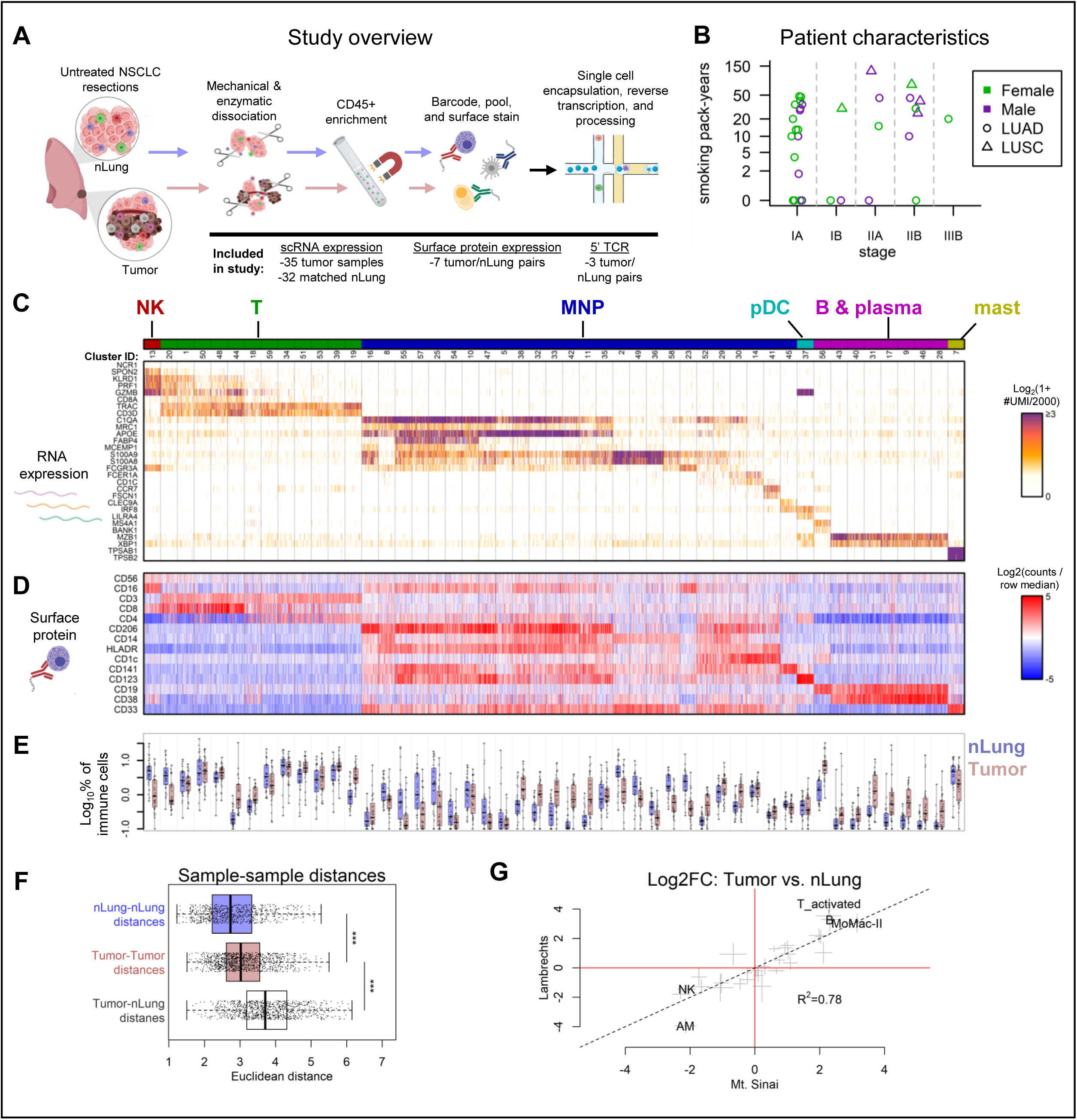
scRNA- and CITE-seq establish the diversity of transcriptional states in the tumor microenvironment. **A**, Study overview. Resected specimens of tumor tissue and non-involved lung (nLung) were digested to single cell suspensions, enriched for CD45+ cells, and subjected to single cell assays including CITEseq and TCRseq. **B**, Clinical data of patients undergoing resection indicating summary pathological stage, smoking history, histological diagnosis, and sex. **C**, Expression of cell type marker genes across scRNAseq clusters of immune cells, grouped by lineage annotation (MNP: mononuclear phagocyte; pDC: plasmacytoid dendritic cell). Heatmap shows the number of unique molecular identifiers (UMI) per cell. Clusters are shown using an even number of randomly selected cells from 7 matched tumor and nLung sample pairs who were analyzed by CITEseq. Cells were downsampled to 2000 UMI/cell. **D**, Expression of lineage-defining surface markers on single cells, as measured by CITEseq. Single cells correspond directly to cells shown in (**C**). CITEseq count values were first quantile normalized across patients, then row-normalized across cells in the heatmap. **E**, Cells per cluster as a percent of total immune cells across 35 tumor and 32 matched nLung samples. Clusters correspond directly to those shown in (**C**) and (**D**). **F**, Box-plots of Euclidean distances between pairs of samples among nLung only (nLung-nLung), tumor only (tumor-tumor), or between nLung samples and tumor samples (Tumor-nLung). Distances between pairs of patient-matched samples were excluded from the Tumor-nLung distribution to prevent confounding due to patient-specific effects. *** P < 0.001, Wilcoxon rank-sum test. **G**, Log-ratios between cell type frequencies in tumor and nLung. Clusters were grouped by cell type annotation. Crosses represent error bars showing the mean ± SEM of Log_2_FC estimates of differences in cell type frequency between tumor and nLung using the cohort collected in the present study (Mount Sinai; x-axis) or the cohort in ref. ^5^ (y-axis).

ScRNA profiles in tumor and nLung were clustered together using a batch-aware algorithm we recently developed in order to combine data across patients, while accounting for batch-specific background noise^16^ (Figures 1C and S1A-C). To develop a general gene-expression model of clusters representing different cell types and states, we relied only on 19 nLung and 22 tumor samples processed with 10X Chromium 3’ V2 chemistry. We then used this model to classify cells from additional samples processed with different protocols or from different datasets showing similar transcriptional profiles. (Figure S1B-D, methods). The RNA-based clustering identified 49 immune clusters within 6 compartments including subsets of T cells, B cells, plasma cells, mast cells, plasmacytoid dendritic cells (pDC), and mononuclear phagocytes (MNP) consisting of macrophages (MΦ), monocytes, putative monocyte-derived dendritic cells (MoDC), and conventional dendritic cells (cDC; Figure 1C and S1E, F). Overall, 377,549 single cells from 35 tumors and 32 matched nLung samples from patients at Mount Sinai were classified into 6 compartments and 30 annotated transcriptional states. CITEseq data further confirmed cell identities using well-established protein markers (Figure 1D). For example, annotation of pDC was based on expression of transcripts associated with this lineage (*LILRA4, IRF8*; Figure 1C) and high expression of known population-defining surface markers (CD123; Figure 1D). While cluster frequencies varied widely among patients, clusters mapped between 590 and 23812 cells, and all clusters included cells from multiple patients (Figure 1E and S1F).

We first compared the variability of samples from different regions within individual tumors to the variability between different patients’ tumors with respect to immune cell type composition. To do this, we examined samples from a study that analyzed three regions per tumor in 8 patients^5^, mapping cells to the clusters produced with our expanded dataset. Clustering the samples by correlation of cell type frequencies among immune cells demonstrated that samples almost always clustered by patient (Figure S1G), and similarly, the Euclidean distances between patient-matched samples of different tumor regions was strongly reduced compared to the distances between samples from different patients (Figure S1H). Therefore, while the total level of immune content may still vary regionally in and around tumors^17^, these analyses demonstrated that inter-tumor differences drive lung tumor immune variability in terms of the phenotypic makeup of the immune cells that are present.

To understand whether the immune changes between tumor and nLung were distinct across patients or, alternatively, globally similar, we estimated the immune diversity within tumor and nLung using the Euclidean distances between the log-transformed cluster-frequencies. This analysis indicated that nLung samples were significantly more homogeneous (Figure 1F; “nLung-nLung distances”) than tumor samples (“Tumor-Tumor distances”; t=8.3; p<2.2e-16). We further compared distances among nLung and among tumor to the distances between nLung and tumor. This analysis showed that the diversity between independent (unmatched) tumor and nLung samples was larger than the diversity within tumor samples (t=19.6; p<2.2e-16) and nLung samples (t=24.6; p<2.2e-16), suggesting that immune landscapes within the TME were significantly changed compared to non-involved tissues, and that most tumors harbored many conserved changes (Figure 1E-F).

We next sought to test if the differences between nLung and tumor could be observed in an independent cohort. The cell type frequencies of 8 matched tumor-nLung pairs described in ref.^5^ indeed validated the distinct microenvironments we observed in our cohort (Figure 1G). This result demonstrated that the observed tumor signatures were robust and reproducible, encouraging us to further study the transcriptional states within it.

### The intratumoral dendritic cell compartment is characterized by expansion of monocyte-derived DC

We next investigated the heterogeneity with the myeloid compartment, given that different myeloid cells have various important roles in generating or inhibiting tumor directed immune responses, including antigen presentation, T cell co-stimulation, and shaping the cytokine milieu within the TME^13^. We identified conventional DC1 (cDC1) expressing *IRF8, WDFY4*, and *CLEC9A* transcripts (Figure 2A) as well as CD141 and CD26 surface markers (Figure 2B), and cDC2 expressing high *CD1C* and *FCER1A* transcripts as well as CD1c and CD5 protein. We also detected a DC cluster expressing *FSCN1* and *CCR7* transcripts and elevated HLADR, CD86, PD-L1, and CD40 surface protein which we described as mature DC enriched in regulatory molecules (mregDC) in great detail elsewhere^18^; in this study we found mregDC were correlated with tumor-antigen uptake and thus help define antigen-charged DC^18^. This phenotype was also consistent with an activated DC phenotype detected in lung and liver tumors by others^6,19^. We furthermore identified clusters that expressed cDC2 markers such as CD1c and *CLEC10A*, but also expressed high levels of monocyte and MΦ genes including *S100A8, S100A9, C1QA*, and *C1QB*, lacked surface expression of the pre-DC surface marker CD5^20,21^, and exhibited increased expression of CD11b and CD14 (Figure 2A, B); we annotated such clusters as MoDC. Importantly, MoDC were distinct from MΦ based on higher levels of CD1c surface protein in addition to their upregulation of the DC2-like transcriptional signature (Figure 1C, clusters 52, 29, and 30). Overall, MoDC were the most prevalent DC subtype and were increased in tumors compared to nLung, whereas mregDC were the rarest (Figure 2C and S2A-C). As we had seen previously^7^, the fraction of cDC1 were strongly reduced in tumors (Figure 2C).

**Figure 2.**
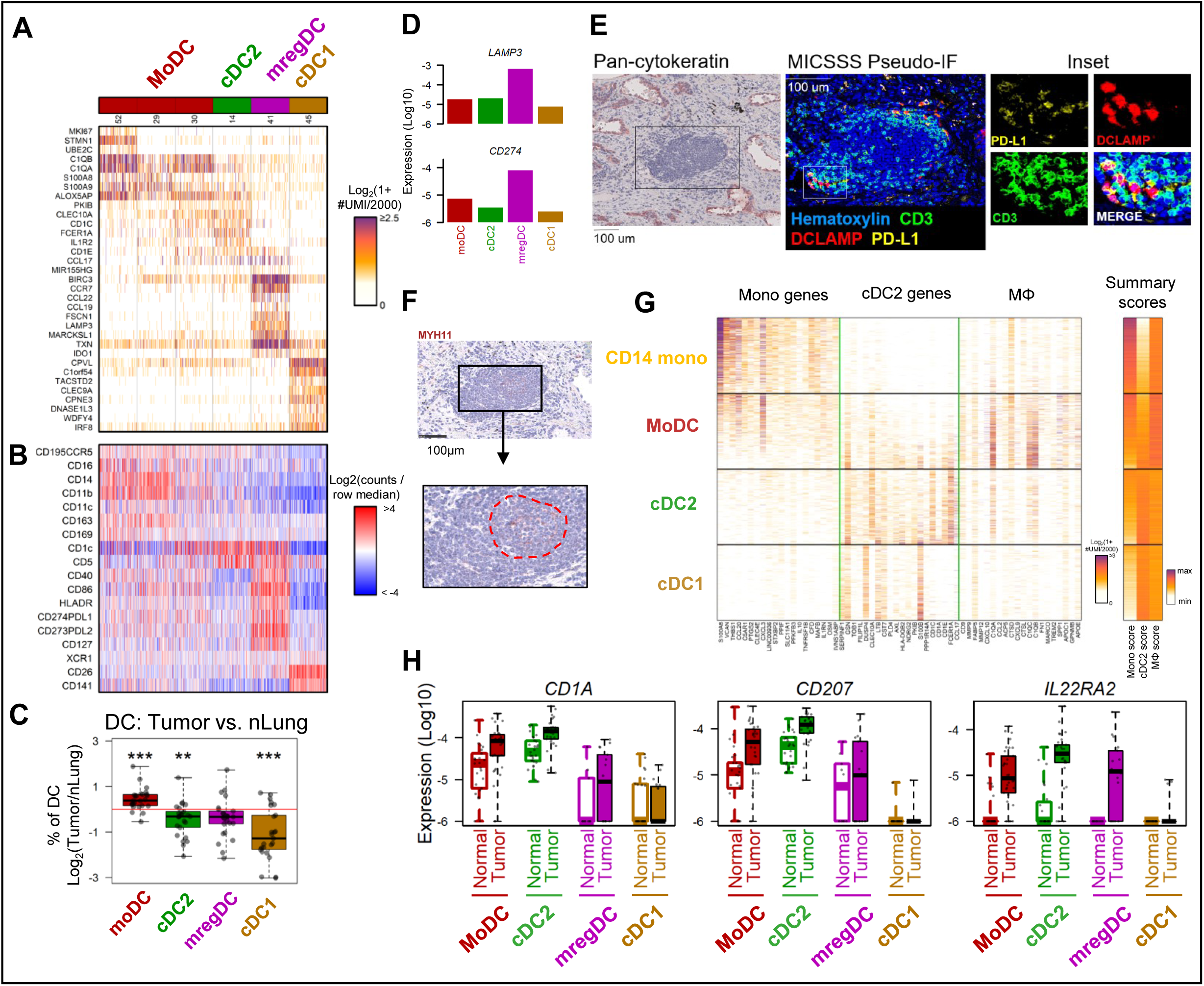
Intratumoral DC comprise expanded MoDC and express an LCH-like signature. **A**, Expression of key genes discriminating scRNAseq clusters of DC, grouped by cell type annotation (MoDC: monocyte-derived DC; cDC: classical DC). Heatmap shows the number of UMI per cell. Clusters are shown using an even number of randomly selected cells from each, drawing from patients who were analyzed by CITEseq with the DC panel shown in (**B**) (4 matched tumor-nLung tissue pairs). Cells were downsampled to 2000 UMI/cell. **B**, Expression of DC surface markers on single cells, as measured by CITEseq. Single cells correspond directly to cells shown in (**A**). CITEseq count values were first quantile normalized across patients, then row-normalized across cells in the heatmap. **C**, Differences between tumor and nLung of DC frequencies normalized to total DC; *P<0.05, **P<0.01, ***P,0.001 (Wilcoxon signed-rank test with Bonferroni correction; N=25 matched tissue pairs with >50 DC observed in each tissue). **D**, Barplots showing average expression of *LAMP3 (DC-LAMP)* and *CD274 (PD-L1)* across DC clusters. **E**, MICSSS imaging showing spatial distribution of DC-LAMP+/PD-L1+ DC in proximity to T cells in a TLS. **F**, Expression of follicular dendritic cell marker MYH11 in TLS in an adjacent section to that shown in (**E**). **G**, Expression among CD14+ monocytes and DC of monocyte, cDC2, and MΦ cell type specific gene signatures (See Figure S2D, E). Heatmaps show expression of 20 genes from each score among single-cells evenly sampled by cell type (left) and as corresponding summary scores. Cells were ordered by the ratio of monocyte:cDC2 summary scores and were downsampled to 2000 UMI. **H**, Boxplots showing average expression of LCH-like signature genes across DC populations in distinct nLung and tumor samples.

Since the activation profile of mregDC is crucial for inducing tumor directed T cell responses^18^, we examined the mregDC distribution in tumors by multiplexed immunohistochemical consecutive staining on a single slide (MICSSS)^22^. We stained for DC-LAMP and PD-L1, as the transcripts of these genes (*LAMP3* and *CD274*, respectively) were highly enriched in the mregDC cluster (Figure 2D). We found that mregDC expressing DC-LAMP and PD-L1 accumulated in tertiary lymphoid structures (TLS) in close proximity to T cells (Figure 2E). CD3-negative areas of TLS, which are putatively analogous to lymph node B cell zones, were frequently populated by MYH11+ follicular dendritic cells^23^, a stromal cell type commonly found in B cell zones (Figure 2F).

To better understand the relationship between MoDC and other MNP, we searched for genes that were mutually exclusive among CD14 monocytes, cDC2, and MΦ (Figure S2D, Table S4). Scoring MoDC using these gene lists in comparison with other MNP populations revealed that MoDC were distinct from MΦ and CD14+ monocytes. Ordering cells within each of these compartments by the expression of these distinct monocyte- and cDC2 gene programs demonstrated anticorrelation of these gene sets among MoDC but not cDC2 (Figure 2G; rho= -0.33, p<2.2e-16; rho=0.016, p=0.29, respectively), demonstrating that MoDC inhabited a phenotypic spectrum between monocytes and cDC2-like cells. While some MΦ genes were expressed in MoDC higher than in cDC2 cells, MoDC were distinct from MΦ based on higher cDC2 gene expression and lower MΦ gene expression (score distributions are detailed in Figure S2E).

To further uncover transcriptional programs that were variable among MoDC and cDC without relying on specific cell classifications, we analyzed the covariance structure of variable genes among all DC. This approach resulted in distinct sets of co-expressed genes (gene “modules”) that varied together across cells, independent of cluster assignments (Figure S2F, G, Table S5). Gene module analysis across the DC compartment revealed upregulation in tumors of multiple modules that were mainly restricted to MoDC and DC2 (Figure S2H, I). The gene modules most upregulated in tumors compared to nlung included genes associated with glycolysis (mod39) and cell cycle (mod38), which were mainly expressed in MoDC cluster 52 (Figure S2G-I). Frequent upregulation of many monocyte- or MΦ-like modules (7, 3, 4, 6, 5, 37, 10) was consistent with a higher frequency of MoDC compared to cDC in tumors.

We also identified a cDC2 module (mod34) which was enriched in tumor lesions compared to nLung (Figure S2G, H) and included *CD1A* and *CD207*. These genes mark the lesional cells of Langerhans cell histiocytosis (LCH), a myeloid inflammatory condition driven by enhanced ERK activation^24^; we therefore referred to this module as “LCH-like”. LCH cells produce many inflammatory cytokines that promote the accumulation of Tregs and activated T cells in LCH lesions^25^. Interestingly, *IL22RA2*, encoding the IL22 decoy receptor IL22-BP, was also included in this module (Table S5). IL22 modulates epithelial cell growth and plays a role in tissue protection through modulation of tissue inflammation and in promoting tumor growth through induction of tissue repair^26^. Expression of the IL22 receptor (*IL22RA1*), meanwhile, negatively correlated with survival in KRAS-mutated lung cancer lesions^27^. These genes were mainly induced in the *bona fide* cDC2 cluster, but were also upregulated in MoDC (Figure 2H). Probing DC expression in an independent scRNAseq dataset of NSCLC immune cells^5^ confirmed upregulation of these genes in tumor associated DC transcriptomes (Figure S2J).

### Tumors are dominated by monocyte-derived MΦ that are distinct from alveolar MΦ

While previous studies have demonstrated phenotypic differences between MΦ populating nLung versus tumors^5,7^, they have been limited in their ability to parse specific MΦ subpopulations with potentially distinct ontogeny and function. Our data showed remarkable heterogeneity within the MΦ compartment as demonstrated by the varying expression of classical marker genes among clusters (Figures 3A and S3A). This level of resolution allowed the identification of alveolar MΦ (AMΦ) clusters expressing *SERPINA1* and *PPARG* and a cluster expressing genes consistent with interstitial MΦ (IMΦ), which thus far have only been defined to a limited extent in humans^28,29^. In contrast to AMΦ that self-renew locally independent of blood precursors^30^, IMΦ are thought to be maintained by circulating monocyte pools even in steady state, albeit at lower rates of turnover than in settings of overt inflammation. IMΦ lacked *PPARG* and expressed *MAF* family transcription factors, *MERTK, CSF1R, LYVE1*, and *CX3CR1*^31^. CD14+ and CD16+ monocytes were defined by the expression of *CD14* or *FCGR3A* respectively and the lack of MΦ markers *MRC1, VSIG4*, and *SIGLEC1*. Other MΦ clusters expressed genes such as *MAFB, CEBPD, FCGR2B*, and *CSF1R*, which are indicative of monocyte origin and shared by monocytes and IMΦ; therefore, these clusters were annotated as MoMΦ. A remaining population of MΦ expressed genes consistent with primary granule formation (*AZU1, ELANE, CTSG*) but distinct from bone marrow progenitors due to lack of *MPO*, and also lacked elevation of neutrophil marker genes^6^ *CSF3R, LRG1, FFAR4*, and *VASP* compared to other myeloid cells (Figure S3A). This cluster was referred to as *AZU1+* MΦ.

**Figure 3.**
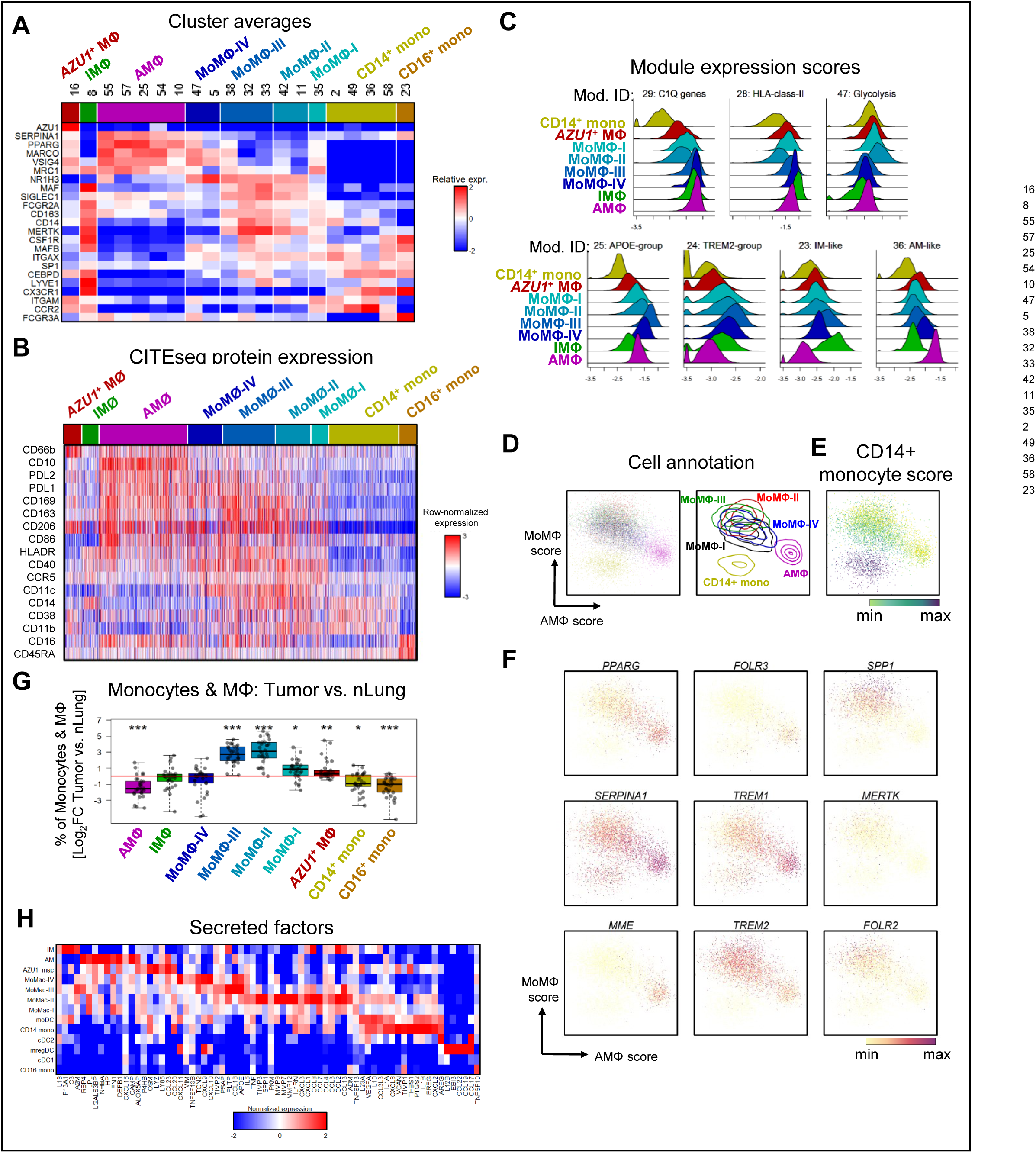
Tumors exclude AMΦ and exhibit a diversity of MoMΦ populations. **A**, Average cluster expression of lineage-defining monocyte and MΦ clusters based on literature review, grouped by cell type annotation. **B**, Expression of myeloid surface markers on single cells, as measured by CITEseq. Clusters are shown using an even number of randomly selected cells from each, from patients who were analyzed with the panel shown (4 matched tumor-nLung pairs). CITEseq count values were first quantile normalized across patients, then row-normalized across cells in the heatmap. **C**, Histograms of gene module scores per cell type (see also Figure S3D-J). **D**-**F**, Expression among CD14+ monocytes, MoMΦ, and AMΦ of cell type specific gene scores. Gene scores were generated based on sets of mutually exclusive, differentially expressed genes among AMΦ, MoMΦ, and CD14+ monocytes (see Figure S3I). Cells are plotted by AMΦ and MoMΦ score, and cell-annotations are indicated by colored dots or contour plots (**D**). Cells are plotted on similar axes and colored by CD14+ monocyte score (**E**), or by expression of individual genes (**F**). **G**, Differences between tumor and nLung of lineage-normalized monocyte and MΦ frequencies; *P<0.05; **P<0.01, ***P,0.001 (Wilcoxon signed-rank test with Bonferroni correction; N=32 matched tissue pairs). **H**, Average cell type expression of secreted factors across MNP cell types.

Using a CITEseq panel of established immune surface markers, we validated the transcription-based cluster annotations and associated new surface markers with the MΦ subpopulations. For example, we found that CD10, not previously appreciated as a MΦ marker, could distinguish AMΦ from other lung myeloid populations (Figures 3B and S3B). This staining was consistent with RNA expression patterns (Figure S3A) and was verified by immunohistochemical staining (IHC) of airspace-residing AMΦ in nLung (Figure S3C). MoMΦ expressed higher levels of CD11c and CD14 than other MΦ populations, whereas IMΦ were notably CD14+/HLADR+/CD11c^int^/CD86-/CD10– (Figure 3B). Thus, CITEseq protein staining confirmed the main MΦ subpopulations identified by the transcriptional classification and defined potential sorting strategies (Figure S3B).

Gene module analysis across all monocyte and MΦ clusters (Figure S3D-G) revealed three broad signatures, consistent with genes that were highly expressed in both AMΦ and MoMΦ (module group I), MoMΦ or IMΦ (module group II), and monocytes (module group III; Figure S3D). Individual modules could be identifiably associated with cell type annotations as well as cell states reflecting, for example, interferon response (modules 32 and 19), heat shock genes (module 49) cell cycle (module 42), HLA class-II expression (module 28), and glycolysis (module 47; Figure S3E). Examining the expression patterns of specific modules across MoMΦ clusters led us to divide them into MoMΦ subtypes I-IV: MoMΦ-II clusters expressed the highest levels of the tumor-enriched module 48, which was driven mainly by *SPP1* and also included IL-1 receptor antagonist *IL1RN*, and module 47 consisting of genes indicating a glycolytically active state (*GAPDH, ENO1, LDHA, ALDOA, TPI1*), and lower levels than other MΦ of *C1Q* and HLA-class-II transcripts (Figures 3C and S3E, G). MoMΦ-III clusters were enriched in module 24 (including *TREM2* and *LILRB4*) and module 25 (including *APOE* and *GPNMB*). MoMΦ-I and MoMΦ-IV were less distinctive than the other MoMac subtypes, but each comprised their own unique gene expression patterns. For example, MoMac-IV expressed the highest levels of module 27 which included *CTSS, CFD*, and *ALDH1A1* and also expressed some genes otherwise confined to AMΦ (module 36), whereas MoMΦ-I was enriched in module 20 (including chemokine ligands *CCL13* and *CCL2*) while MoMac IV was not. Together, these analyses identify multiple tumor MoMΦ phenotypes with distinct metabolic and immunomodulatory gene programs that are enriched in the tumor milieu and likely contribute to defining the tumor microenvironment.

Gene set scores based on mutually exclusive, differentially expressed genes among CD14+ monocytes, AMΦ, and MoMΦ (Figure S3H and Table S4) showed that AMΦ and MoMΦ were each distinct from CD14+ monocytes (Figure 3D) but that MoMΦ expressed a gradient of the CD14+ monocyte score (Figure 3E). Analysis of the gene expression patterns of hundreds of genes within the scores supported the general trends (Figure 3F). MoMΦ clusters were also distinct from the IMΦ cluster based on many transcripts and surface proteins (Figure 3A, B), although some MoMΦ, especially those that were the most distant from AMΦ, shared some IMΦ genes such as *CSF1R, FOLR2*, and *MERTK* (Figures 3A, F and S3A).

The predominant populations that increased in tumors were MoMΦ, while AMΦ were strongly depleted from tumors and IMΦ frequencies were unchanged (Figure 3G). Monocyte frequencies were also decreased, possibly reflecting their differentiation to MoMΦ or MoDC. Given that individual MoMΦ-subsets changed between nLung and tumor to different extents, we asked how these differences related to the underlying phenotypic heterogeneity within the MoMΦ compartment, beyond signatures revealed by module analysis. Selecting for a set of highly expressed transcripts encoding secreted factors demonstrated strong differences between MNP subsets (Figure 3H). MoMΦ-II, the most tumor-enriched subset, expressed the highest levels of inflammatory cytokines *TNF* and *IL6*, transcripts encoding the pleiotropic factor SPP1, a broad collection of matrix metallopeptidases *MMP-7, -9*, and *-12*, as well as CCR2/5 ligands *CCL-2, -8*, and *-7*. By comparison, other MoMΦ populations expressed less distinct secretory profiles. Multiple MNP populations expressed the CXCR3-ligand chemotactic factors *CXCL-9,-10,-11* including MoMΦ, MoDC, and mregDC, while these ligands were distinctly absent from AMΦ, IMΦ, AZU1+ MΦ, monocytes, cDC1, and cDC2. MregDC, meanwhile, expressed distinct cytokines and chemotactic factors associated with T cell engagement, including *IL12B, EBI3, CCL17, CCL22*, and *CCL19*, which were expressed to a minimal or greatly reduced degree in monocytes, MΦ, or MoDC.

### TCRs limited to tumors mark T cells with distinct phenotypic features

CITEseq characterization of T cells identified populations of CD8+ cells that were characterized by an NK-like signature (T_NK-like_), high expression of *GZMK* (T_GZMK_), expression of genes related to tissue-residence such as *ITGA1* transcript and CD103 and CD69 protein (CD8+ T_rm_), and a cluster consistent with activated T cells, expressing high levels of *IFNG, GZMB, LAG3, CXCL13*, and *HAVCR2* transcripts, as well as high PD-1, ICOS, and CD39 protein (T_activated_; Figure 4A, B). Other clusters, which mostly consisted of CD4+ cells, could be separated into T_reg_, T_rm_, cells expressing a profile consistent with either central memory or naïve cells (T_CM/Naïve-like-I_; *TCF7, SELL, LEF1, MAL*, and surface expression of CD127), and a group of clusters expressing both intermediate levels of this signature as well as a tissue-residency signature (T_CM/Naïve-like-II_). Cells within the T_CM/Naïve-like_ clusters did not otherwise segregate by signatures related to antigen experience, TCR engagement, activation, or exhaustion state.

**Figure 4.**
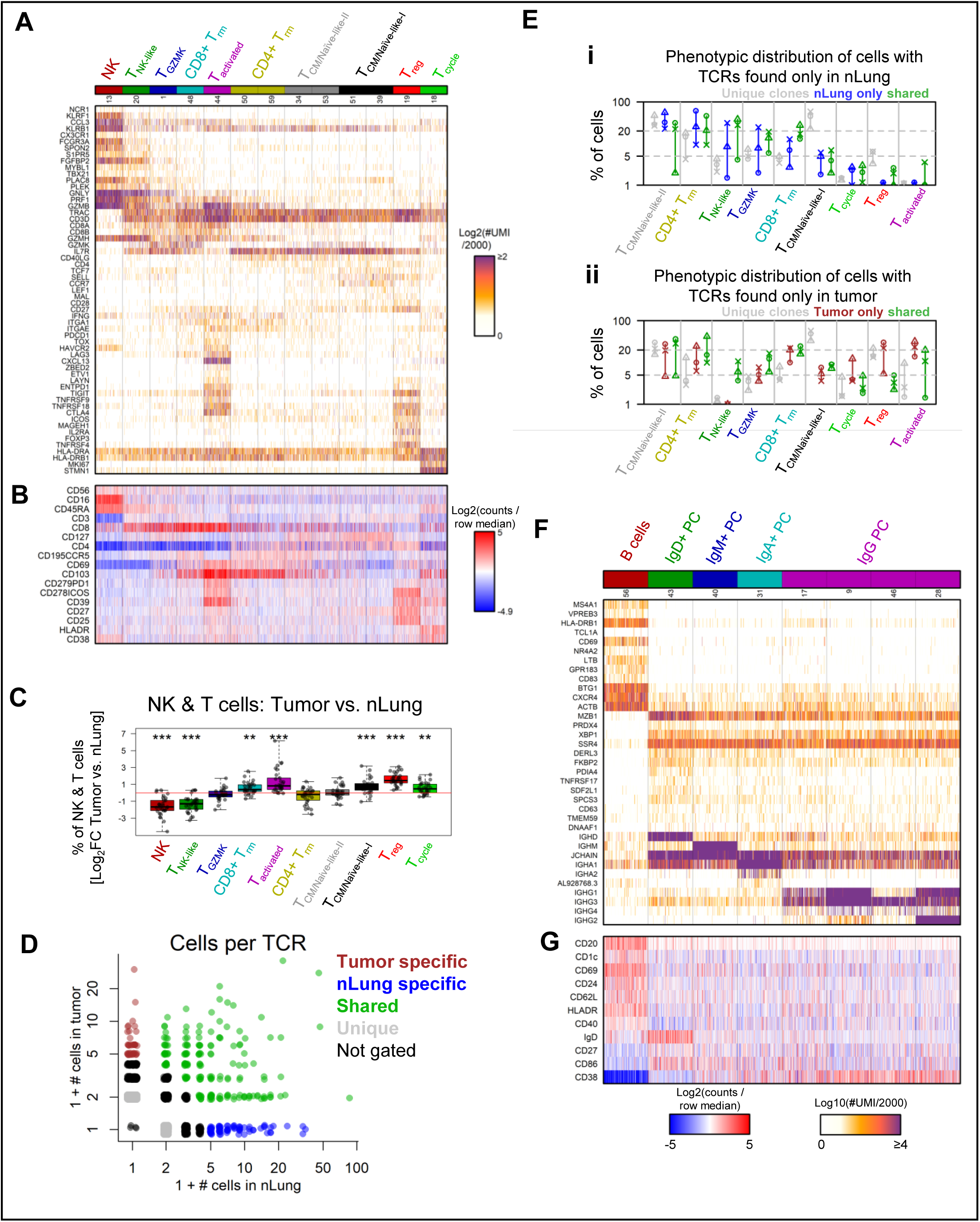
CITEseq and TCR analysis of the adaptive immune compartment. **A**, Expression of key genes discriminating scRNAseq clusters of T cells, grouped by cell type annotation. Heatmap shows the number of UMI per cell. Clusters are shown using an even number of randomly selected cells from each, drawing from patients who were analyzed by CITEseq with the T cell panel shown in (**B)** (2 matched tumor-nLung tissue pairs). Cells were downsampled to 2000 UMI/cell. **B**, Expression of T cell surface markers on single cells, as measured by CITEseq. Single cells correspond directly to cells shown in (**A**). CITEseq count values were first quantile normalized across patients, then row-normalized across cells in the heatmap. **C**, Differences between tumor and nLung of population frequencies normalized by total NK and T cells; *P<0.05; **P<0.01, ***P<0.001 (Wilcoxon signed-rank test with Bonferroni correction, N=32 matched tissue pairs). **D, E**, Phenotypic distribution of T cells among tissue-stratified clonotypes. Frequencies of unique TCRs observed by scTCRseq in nLung (x-axis) or tumor in a representative patient (**D**). In (**E**), cells were first grouped by TCR tissue tropism categories as defined in (**D**); for 3 patients, the phenotypic makeup of the cells with unique TCRs, tissue-specific TCRs, or TCRs shared across tissues is plotted for nLung (**i**) and tumor tissues (**ii**) is plotted as a percent of cells with similarly tissue-distributed TCRs. Each patient is indicated by shape. **F**, Expression of key genes discriminating scRNAseq clusters of B and plasma cells, grouped by cell type annotation. Heatmap shows the number of UMI per cell. Clusters are shown using an even number of randomly selected cells from each, drawing from patients who were analyzed by CITEseq with the B cell panel shown in (**G**) (4 matched tumor-nLung tissue pairs). Cells were downsampled to 2000 UMI/cell. **G**, Expression of B and plasma cell surface markers on single cells, as measured by CITEseq. Single cells correspond directly to cells shown in (**F**). CITEseq count values were first quantile normalized across patients, then row-normalized across cells in the heatmap.

While clustering cells using their transcriptional profiles did not result in complete separation of CD4+ and CD8+ cells, CITEseq allowed for the comparison of CD4+ versus CD8+ cells within otherwise transcriptionally similar groups. The T_activated_ cluster could therefore be separated into CD4+ and CD8+ components (15.5% and 74.8%, respectively). Differential expression analysis between these subsets showed that, on average, CD4+ cells in this cluster expressed increased levels of *CXCL13, CD40LG, BCL6*, and *IL21* (Figure S4A) consistent with a phenotype similar to T-follicular-helper. We next asked whether we could use profiles of CD8+ and CD4+ cells within this cluster to classify CD4 and CD8 cells from samples lacking CITEseq surface staining, which was not available for the majority of our dataset. Learning a signature-based classifier from a training set consisting of the T_activated_ cells from 2 patients and testing this signature on the remaining patients with CITEseq staining demonstrated that transcriptional based classification guided by antibody signals was highly accurate (86% on test set; Figure S4B). This classification could further discriminate cells that uniquely expressed *CD8*-*A/B* transcripts or *CD4* transcripts across the remaining cells in the dataset (84% accuracy; Figure S4C). Applying this classification generally allowed for the separation of CD4 T_activated_ from CD8 T_activated_ across the dataset (Figure S4D). Similar to a recent report^32^, independent quantification of these cells separately and comparing their frequencies demonstrated a high correlation across tumors (Figure S4E; rho=0.58, p=2.7e-4), so they were continued to be grouped for further analysis.

While T_activated_ and T_reg_ were the most increased T cell populations in tumors compared to nLung (Figure 4C), another cluster, characterized by high expression of cell-cycle genes *MKI67* and *STMN1*, and surface expression of HLA-DR and CD38, was also significantly increased in tumors (T_cycle_; Figure 4A-C). Other than expressing these hallmarks of proliferation, the T_cycle_ cluster was diverse with respect to RNA and protein expression (Figure 4A, B). Analyzing the cells comprising T_cycle_ by gene scores constructed from genes differentially expressed among the other clusters demonstrated that T_cycle_ is a mixture of multiple T cell phenotypes that share the cycling state (Figure S4F and Table S4). While tumors expressed overall higher frequencies of cycling T cells (Figures 4C and S4G), T_activated_ and T_reg_ showed the highest frequencies of cycling cells compared to other phenotypes (Figure S4H).

To understand the clonal relationships among T cell phenotypes in tumor and nLung tissues, we performed paired scRNAseq and TCRseq using a nested PCR approach on paired tissues from 3 patients. Classification of the transcriptomes among the clusters and analysis of the T cell repertoires among these phenotypes confirmed that cells mapping to the T_activated_ cluster were the most clonal population in tumors (Figure S4I). Furthermore, dividing clones into groups based on their expansion in nLung or tumor determined the phenotypes of shared clones compared to clones detected in either tissue specifically (Figures 4D, E, S4J, K). In nLung samples, the phenotypic distribution of T cells with TCRs either shared with tumor samples or only present only in nLung was similar (Figure 4Ei). In tumors, however, we observed differences in phenotypic distributions between cells with shared versus tissue-specific TCRs (Figure 4Eii). Specifically, no T_NK-like_ cells in tumors had TCRs that were uniquely expanded in tumors. Furthermore, the proportion of T_cycle_, T_reg_, and T_activated_ among cells with TCRs uniquely expanded in tumors were all markedly increased compared to their proportions among cells with TCRs present in both tissues, and these relationships were not observed in nLung. By controlling for the distribution of cells with shared TCRs in the tumor, we found that the clonal enrichment in these populations was not simply due to enrichment of these phenotypes within tumor. Together, the finding that T_activated_ are enriched in clonally expanded and cycling T cells at the tumor suggests that their accumulation is at least in part due to local clonal expansion.

### B cells and plasma cells are increased in tumors, but the B:plasma cell ratio is conserved between tumor and nLung

B and plasma cells represented the most globally increased lineage among immune cells in tumors compared to nLung across multiple datasets (Figure 1C-E); B cells were increased as a proportion of immune cells by a median of 6.4-fold (IQR: 2.5-8.4), while plasma cells were similarly increased by a median of 4.1-fold (IQR: 2.2-9.4). B cells and plasma cells were strongly distinct both on the RNA and surface marker level (Figure 4F, G). Plasma cell clusters included rare IgD+ plasma cells, which were also the only CD38^int^ population. B cell frequencies overwhelmed plasma cell frequencies with IgD+ and IgM+ plasma cells being the rarest, but lineage-normalized frequencies were not different between nLung and Tumor (Figures 1E and S4L). B cells and plasma cells were therefore found to increase in tumors without significant overall perturbation of the B:plasma cell ratio or plasma cell isotype ratios.

### Ligand-receptor interactions identify potential drivers of an adaptive activation module

In order to identify links between cellular phenotypes that may drive patient diversity, we performed correlation analyses across cell type frequencies in tumors normalized within lineage (Figure 5A). Among the most highly correlated cell types were T_activated_, IgG+ plasma cells, and MoMΦ-II; we therefore called these cell types collectively the lung cancer activation module (LCAM). The cell types that were most anticorrelated to this module included B cells, T_cm/naïve-II_, AMΦ, resting cDC, and AZU1+ MΦ. Sorting patients by these cell types revealed that patients could be broadly grouped into those with high or low frequencies of LCAM cell types (Figure 5B). We called these groups LCAM^hi^ and LCAM^lo^, respectively. Including samples from external datasets in this stratification supported the overall pattern (Figure 5B and S5A). This stratification was not strongly associated with changes in lineage frequencies among total immune cells, and accordingly, samples from both LCAM^hi^ and LCAM^lo^ groups generally displayed lineage-population shifts in line with overall tumor versus nLung differences, such as decreased NK and increased B lineage frequencies (Figure 5C). Therefore, while LCAM cell types included some of the populations that were most enriched in tumor compared to nLung on average, the LCAM axis was importantly not a reflection of tumor sample purity.

**Figure 5.**
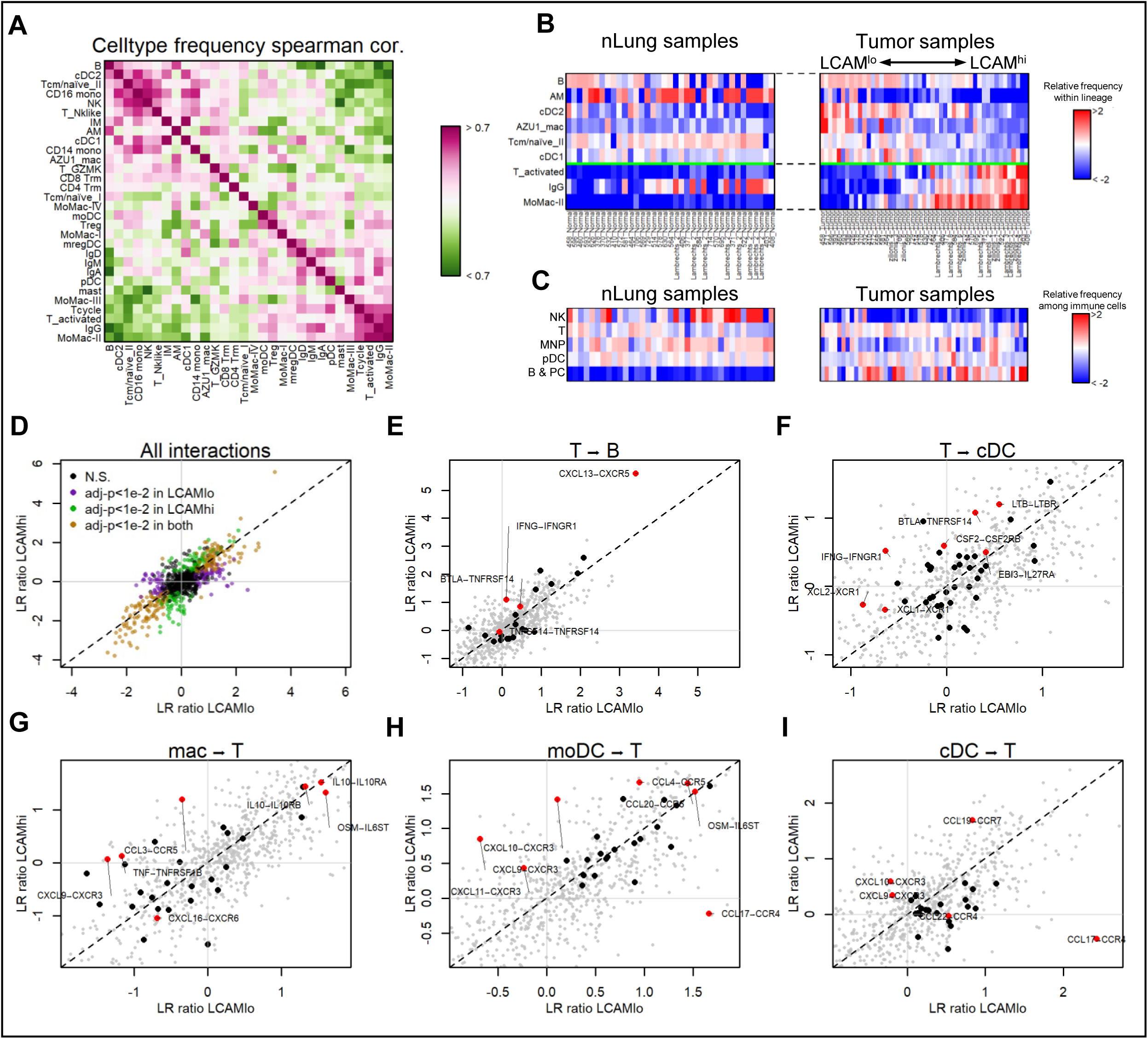
Cell-cell interactions drive an axis of adaptive activation. **A**, Spearman correlation of cell type frequencies after normalization within lineage. Analysis includes 23 tumors that were processed similarly using 10X Chromium V2 and CD45+ magnetic bead enrichment. **B**, Lineage-normalized cell type frequencies of LCAM^hi^ and LCAM^lo^ cell types among pooled nLung and Tumor samples from Mount Sinai and refs. (^5,6^) (50 tumor patients with 40 matched nLung samples). nLung samples are ordered to match the order of tumor samples based on frequencies of LCAM celltypes. **C**, Immune lineage frequencies of nLung and Tumor samples; with columns corresponding to patient ordering in (**B**). **D-I**, Log2 Ratio of ligand-receptor (LR) intensity scores between tumor and nLung of LCAM^hi^patients, (“LR ratio”; y-axis) and LCAM^lo^ patients (x-axis). All interactions among T cells, B cells, MΦ, MoDC, cDC, and monocytes, colored by indication of significance (permutation test, **D**). Dashed diagonal line indicates unity. **E-I**, Showing same data as in (**D**), but highlighting in bold LR ratios for interactions between T cell ligands and B cell receptors (**E**), T cell ligands and cDC receptors (**F**), MΦ ligands and T cell receptors (**G**), MoDC ligands and T cell receptors (**H**), and cDC ligands and T cell receptors (**I**). Labelled interactions are plotted in red.

To identify tumor-specific immune dysregulation that may contribute to shaping the LCAM^hi^ vs. LCAM^lo^ cellular organization among patients, we performed an unbiased analysis of ligand-receptor pairs between immune subsets, leveraging a dataset of secreted ligands and their experimentally validated receptors^16,33^ (Figure S5B, C and Table S6), comparing differences in ligand-receptor (LR) intensity scores^16^ between LCAM^hi^ and LCAM^lo^ groups, as well as between each group and their respective adjacent nLung tissues (Table S7). Overall, both LCAM^hi^ and LCAM^lo^ patients demonstrated correlated modes of LR activation in tumors compared to nLung (Figure 5D). In particular, tumors in both LCAM^hi^ patients and LCAM^lo^ patients exhibited strong intensity scores between T-cell derived CXCL13 and B cell CXCR5, which is likely contributing to the influx of B and plasma cells seen in tumors (Figure 5E). T cells in LCAM^hi^ but not LCAM^lo^ patients also produced other factors in tumors but not nLung capable of stimulating B cells through the IFNG-IFNGR1 axis and the BTLA-TNFRSF14 axis (Figure 5E). In addition, B cells from LCAM^hi^ but not LCAM^lo^ patients highly expressed TNFSF9 (41BBL), which ligates TNFRSF9 (41BB), that we found highly expressed on T_activated_ cells (Figure S5D), indicating B cells from LCAM^hi^ patients participate in activation of T cells via TNFSF9-TNFRSF9 interaction.

In addition, we observed increased IFNG-IFNGR signaling between T cells and myeloid cells in LCAM^hi^ patients (Figures 5F and S5E, F). Potentially in result, LCAM^hi^ patients displayed higher activation of the CXCL9/10/11-CXCR3 axis between myeloid and T cells (Figure 5G-I). Whereas MΦ and MoDC demonstrated many conserved ligands upregulated in both LCAM^hi^ and LCAM^lo^ tumors such as IL10 and OSM (Figure 5G, H; note distribution of highlighted LR pairs along the diagonal), tumor cDC shared few ligands between the two groups, and rather upregulated CCL19 higher in LCAM^hi^ patients compared to CCL17 in LCAM^lo^ patients (Figure 5I); MoDC also demonstrated the latter pattern, selectively expressing CCL17 in LCAM^lo^ patients (Figure 5H), suggesting that DC expression of CCL19 may be a unique feature of cDC necessary for activation of T cells as well as induction of a humoral immune response. Overall, differences in ligand-receptor intensity scores between LCAM^hi^ and LCAM^lo^ patients supported such a patient stratification, and provide possible mechanistic insight into immune-cell crosstalk underlying the development of the LCAM axis, including IFNg signaling as a major driver.

### Projection of bulk-transcriptomic data onto scRNA-derived signatures reveals the presence of the LCAM^hi^ module in two independent LUAD datasets

To identify tumor-related correlates of the LCAM module, we aimed to analyze a larger patient cohort in order to increase statistical power. Therefore, we implemented an unbiased method of scoring bulk transcriptomic signatures along the LCAM axis^16,34^. Specifically, we identified genes that were both differentially expressed between LCAM^hi^ and LCAM^lo^ tumor samples (Figure S6A), and also highly specific to the cell types enriched or depleted in the LCAM^hi^ tumors (Figure S6B, see methods). Using a published tabulation on estimated immune content of 512 lung adenocarcinoma (LUAD) patients available from TCGA based on expression of immune genes of all lineages^35^, we saw as expected that scores generated with either gene set were highly correlated with estimates of overall immune content (Figure S6C, D), but an ensemble LCAM score computed by the difference of these scores (LCAM^hi^ score – LCAM^lo^ score) was not (Figure S6E). As predicted by our scRNAseq data, when controlling for the immune content, we generally observed negative correlations of LCAM^hi^ and LCAM^lo^ gene scores among tumors except for samples with the 10% lowest immune content (Figure S6F, G), suggesting that the ensemble LCAM score might measure a mode of immune activation that is independent of the overall immune infiltration measured by immune content. We excluded the samples with the lowest 10% immune content from further analysis because probing the immune signatures was likely less informative within these samples. Sorting the patients by the ensemble LCAM score revealed the presence of LCAM^hi^ and LCAM^lo^ patient groups within the cohort (Figure 6A).

**Figure 6.**
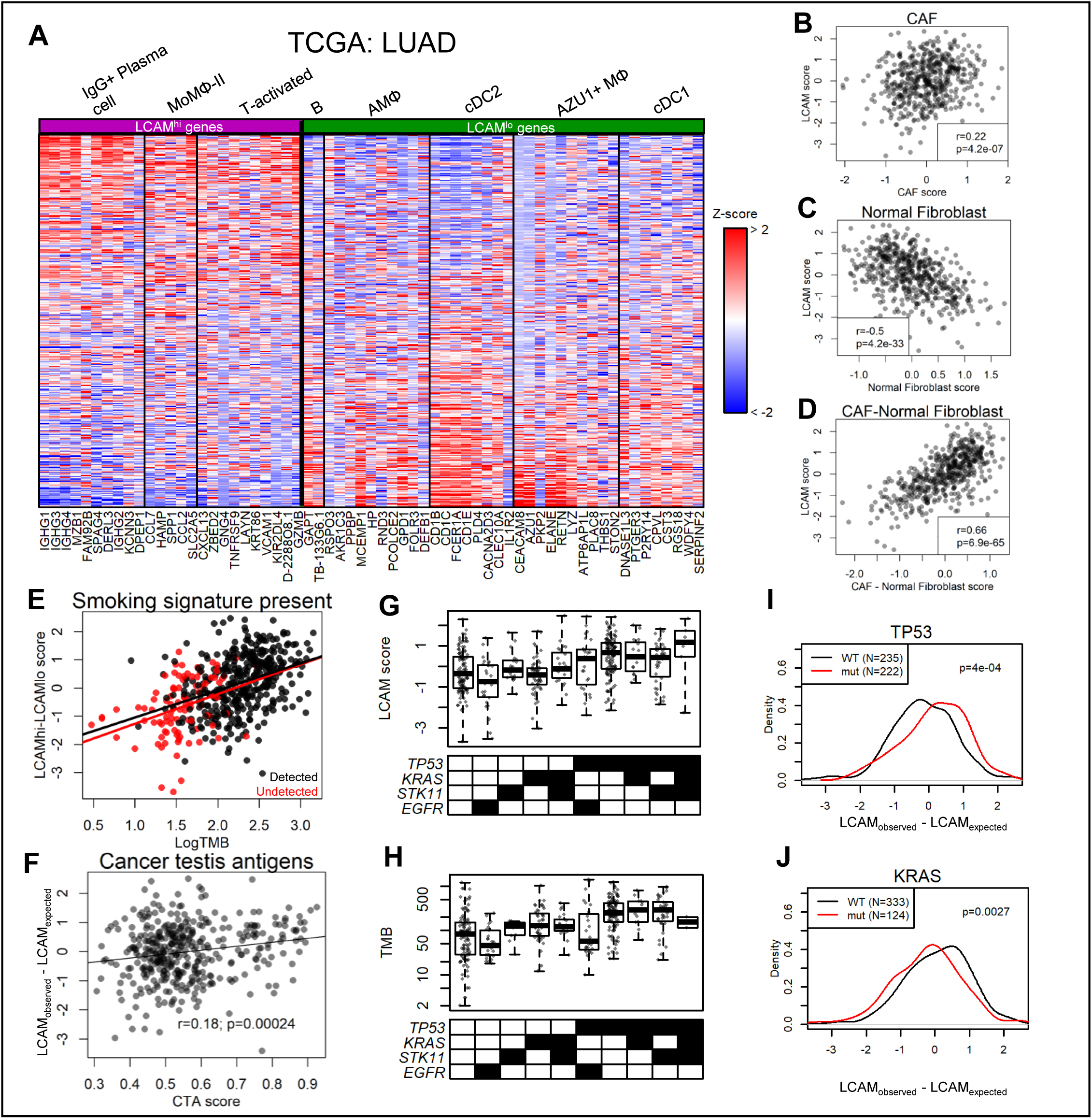
Tumor features related to the LCAM immune response. **A**, Normalized expression of LCAM^hi^ and LCAM^lo^ bulk-RNA signature genes, determined as shown in Figure S6A, B and as described in the methods, in TCGA lung adenocarcinoma dataset. Cell type association with sets of genes for each signature is shown. Patients are sorted along y-axis by ensemble LCAM score. **B-D**, Scatter plots of the ensemble LCAM score (y-axis) with signature scores based on genes that are specific for CAFs (**B**), normal fibroblasts (**C**), or the difference between these scores (**D**) in TCGA lung adenocarcinoma data. Stromal signatures are based on the stromal data reported in ref. ^5^. **E**, Scatter plot of LogTMB and ensemble LCAM score. Patients are divided into those with presence of a smoking-related mutational signature (black) and those without presence of the signature (red). Black and red lines indicate linear regression relationships computed over each group of patients independently (r=0.38; p=9.2e-5 in the undetected smoking signature group; r=0.34; p=1.1e-12 in the detected signature group). **F**, Scatter plot of Cancer testes antigen expression score (CTA score), as computed in ref. ^53^, and the residuals of the regression of the ensemble LCAM score on the LogTMB. **G and H**, Boxplots showing either the ensemble LCAM score **G**), or TMB (**H**) among TCGA lung adenocarcinoma patients, divided by combinations of driver mutations. **I and J**, Histograms of residuals of the regression of the ensemble LCAM score on the LogTMB, with patients stratified by *TP53* (**I**) or *KRAS* (**J**) mutational status (Two-sided t-test).

To see whether a similar pattern was present in additional datasets, we probed the independent CPTAC: LUAD cohort, consisting of 110 treatment-naive LUAD patients undergoing surgical resection on whom bulk RNAseq and WES had been performed (Michael A. Gillette, et al. *Cell*, In press). Similar to the TCGA cohort, sorting the patients by the ensemble LCAM score revealed the presence of LCAM^hi^ and LCAM^lo^ patient groups (Figure S6H), further establishing the prevalence of this cellular module in a subset of LUAD patients.

### LCAM immune response correlates with tumor-genotype and expression of tumor-antigens in LUAD lesions

While the anti-tumor immune response can be modulated by many tumor-intrinsic and tumor-extrinsic factors, the tumor-infiltrating immune cells exist as part of a complex microenvironment that includes many other stromal populations^13,36^. To ask whether the ensemble LCAM score is associated with other non-tumor, non-immune stromal populations, we derived gene lists that were specific for individual stromal populations identified in a public dataset of 8 NSCLC patients^5^ (Table S4), and used these genes to quantify enrichment of stromal populations in TCGA LUAD data. The ensemble LCAM score correlated with a cancer-associated fibroblast (CAF) enrichment score, anticorrelated with a normal fibroblast enrichment score, and strongly correlated with the difference of these scores (Figure 6B-D). Meanwhile, it exhibited weak or absent correlations with a tumor-associated blood endothelial cell (BEC) enrichment score, an nLung BEC enrichment score, and a lymphatic endothelial score (Figure S6I). These data suggest an intimate link between development of the LCAM cellular module and a CAF-like fibroblast phenotype, which should be explored in further detail as CAF have been suggested to act as major regulators of TIL function^36,37^.

We hypothesized that variability in immune and stromal states captured by the LCAM and CAF signatures could be associated with different tumor properties. While the ensemble LCAM score demonstrated a small but significant increase in large tumors (t=2.60, p=0.01 between TNM T-stage=T1 and T-stage>T1), we observed variable LCAM presence among tumors of all stages (Figure S6J). Furthermore, while PD-L1 expression is the most commonly used biomarker guiding ICB treatment, we also observed a weak correlation between the ensemble LCAM score and total *CD274* expression (r=0.21, p=2.7e-5). TMB, meanwhile, has been demonstrated to be one of the most robust predictors of checkpoint response^38^, and is supported by the key mechanistic hypothesis that tumors with many mutations more easily activate and are targeted by the immune system via the generation of mutated peptides and damage-associated molecular patterns. Strikingly, the data showed that the ensemble LCAM score was strongly correlated with TMB both in TCGA (Figure 6E; r=0.47 p<2.2e-16) and in CPTAC (Figure S6K) (r=0.53 p=2.3e-9). By comparison, other scores measuring overall immune content (Immune ESTIMATE^35^) or specific aspects of immune state (T cell-inflamed gene expression profile (GEP) score^39,40^) had much weaker associations with TMB (Figure S6L, M). Importantly, correlation with TMB was observed broadly across LCAM^hi^ genes expressed in multiple cell types, whereas conversely, anti-correlation with TMB was also observed broadly across LCAM^lo^ genes (Figure S6N). The ensemble LCAM score correlated with TMB to similar extents among patients grouped within each TNM T-stage (Figure S6O).

In LUAD cases, TMB is strongly associated with smoking history. Consistent with this relationship, the ensemble LCAM score correlated with smoking pack-years (rho=0.23, p=4.4e-5). Therefore, smoking history confounded the correlation we observed between TMB and the ensemble LCAM score, suggesting that the immune signature could be only indirectly related to mutations and specifically mutated neoantigens, but rather due to alternate modes of immune dysregulation related to smoking exposure. To test this hypothesis, we stratified tumors by the detection of the smoking-related mutational signature characterized by C>A de-aminations within specific trinucleotide contexts^41,42^. This approach removed uncertainty related to unreliability of patient-reported smoking statistics and missing clinical data. We observed that both tumors with and without detection of this signature exhibited significant correlations between TMB and the ensemble LCAM score (r=0.38; p=9.2e-5 in the undetected smoking signature group) despite having clearly distinct distributions of TMB (Figure 6E), suggesting that this relationship was independent of smoking-driven immunomodulation.

We then asked which additional features of the tumors may influence the ensemble LCAM score beyond the effect caused by differences in TMB. To perform this analysis, we regressed the ensemble LCAM score onto the LogTMB and correlated candidate variables with regression residuals, which quantify the difference between the observed and expected LCAM scores based on this relationship. For example, scores quantifying total predicted single-nucleotide-variant- or Insertion/deletion-induced neoantigens did not correlate with these differences (Figure S6P, Q), indicating that these neoantigen prediction scores did not provide more information regarding the LCAM immune modulation than TMB alone. However, consistent with the hypothesis that generation of tumor-associated antigens was the key mechanism connecting TMB to an LCAM response, we found that a score quantifying total tumor associated but not tumor-specific cancer-testis antigens (CTA) was correlated with the regression residuals (Figure 6F; r=0.16, p=3.4e-3), suggesting that additional tumor-associated antigens beyond those directly caused by tumor mutations may also contribute to induction of the LCAM response.

Most adenocarcinoma patients have at least one of a small number of common driver mutations, including *KRAS, EGFR, STK11*, and *TP53*. Recently, it was shown that LUAD patients responsive to immune checkpoint blockade frequently have tumors harboring *TP53* mutations, and that *TP53* mutant status was associated with enrichment of CD8 T cells in the TME^43,44^. However, immune-related effects of individual mutations have generally not been considered independently given their correlation with TMB. Specifically, while *TP53*-mutant tumors had higher ensemble LCAM scores compared to *TP53*-WT/(*EGFR* or *KRAS* or *STK11*)-mut tumors in both TCGA and CPTAC datasets (Figures 6G and S6R), *TP53* was also most strongly associated with increased TMB (Figures 6H and S6S). In order to statistically test whether these mutations were associated with higher LCAM scores while controlling for TMB, we regressed the LCAM score onto the LogTMB and asked whether any individual mutations were correlated with the regression residuals. Interestingly, this analysis showed that *TP53*-mutant patients had higher LCAM scores than expected by a model assuming only correlation with TMB (Figure 6I; p=1.4e-3). *KRAS*-mutant patients, meanwhile, had lower LCAM scores than expected by this model (Figure 6J; p=1.6e-4). There was no similar deviation seen in either *STK11*- or *EGFR*-mutant patients (Figure S6T). Overall, projection of bulk signatures onto axes defined by variation in our scRNAseq cohort suggested that expression of the LCAM cellular module is a marker of adaptive response against mutated and ectopically-expressed tumor-associated antigens that is independent from the overall level of immune infiltration.

## DISCUSSION

The analysis of matched tumor and nLung tissues from 35 patients as described here provides the largest unbiased single-cell map of the immune response of early-stage lung cancer lesions to date. CITEseq analysis, combining phenotypic classifications based on surface protein expression with transcriptomic profile, serves here to help unite high-dimensional models of cellular classification and refine our understanding of the immune cellular landscape in disease lesions. By further integrating tumor and nLung samples from public datasets, we demonstrated the robustness of the reported signatures across platforms. Importantly, based on high levels of changes conserved across tumor lesions, these data support the notion that common immunotherapy treatment paradigms could be beneficial for large subsets of patients despite existing disease heterogeneity.

Among tumors, patients could, however, be stratified along a dominant LCAM axis that was independent of overall immune infiltration or changes in proportions of immune lineages. This axis was defined by a high level of IgG+ plasma cells, activated T cells that were clonally enriched in the tumor and expressing a proliferation signature, and MoMac-II that expressed *SPP1*, a glycolysis signature, and a set of inflammatory secreted factors; this module of cell types anticorrelated with B cells, T cells with a Tcm/naïve-like phenotype, resting cDC, AMΦ, and MΦ expressing *AZU1*. We therefore propose that LCAM^hi^ patients are undergoing a more vigorous antigen-specific antitumor adaptive immune response, whereas LCAM^lo^ patients fail to mount an adaptive response to such a degree. Unbiased ligand-receptor analyses showed that, while both LCAM^hi^ and LCAM^lo^ tumors expressed similar patterns of ligand-receptor pairs among immune cells compared to nLung, LCAM^hi^ status was specifically related to heightened *CXCL13* expression by T cells, IFNg signaling from T cells to myeloid and B cells, and CXCL-9,10,11 signaling from myeloid cells; cDC meanwhile expressed more *CCL19* in LCAM^hi^ tumors compared to more *CCL17* in LCAM^lo^ tumors. These factors likely served to modify the immune response around a set of conserved changes compared to nLung observed in both LCAM^hi^ and LCAM^lo^ tumors.

When analyzed in the context of broader datasets with paired bulk transcriptomics and whole exome sequencing, an ensemble gene score learned from the LCAM^hi^ and LCAM^lo^ patients and associated cell types strongly correlated with measures indicative of high levels of tumor-associated antigens, namely TMB and a cancer testis antigen score. Interestingly, the LCAM score was not correlated with the overall immune infiltrate, and was independent of the T-cell inflamed gene expression profile score commonly used to reflect immune activation in tumor lesions^39,40^. While the ensemble LCAM score was correlated with smoking status and weakly correlated with stage, the relationship with TMB remained even after controlling for these possible confounders. Given that TMB has demonstrated predictive power with response to ICB response in NSCLC^14,38^, the relationship between TMB and the LCAM cellular module in treatment-naïve patients suggests that this effect may be mediated via a conditioning of the immune system that exists prior to treatment, and that measurement of this cellular module may provide a more direct indicator with respect to the immune system’s propensity for ICB response. Specifically, the fact that many factors significantly influence the LCAM score, not just TMB, demonstrates how the immune system integrates multiple types of signals to establish its set point.

Importantly, while previously reported immune signatures have been proposed to reflect tumor cytolytic activity or T cell and IFNg-driven immune response in association with tumor antigens and immune evasion modes^39,40,45^, the LCAM axis presented here represents an integrated assessment of the immune cellular organization, based on all immune cell types as defined by scRNA across patients, likely arising as a direct response to tumor antigens.

An additional question of clinical interest relates to how different driver mutations affect the conditioning of the immune system and ICB response. The analysis presented here shows that the common LUAD driver mutations *EGFR* and *STK11* had little effect on the LCAM response beyond that explained by their association with TMB. While *STK11* mutation status has been shown to be the most prevalent genomic driver of primary resistance to ICB^44,46^, there were no patients with *STK11*-mutated tumors in our scRNAseq cohort, so this effect can therefore not be addressed here. Meanwhile, compared to what was expected based on each tumor’s TMB alone, *TP53* mutation intensified the LCAM response and *KRAS* mutation blunted it. Interestingly, the latter result is consistent with a recent report demonstrating that pharmacological blockade of KRAS-G12C in preclinical studies resulted in a robust immune response and synergized with anti-PD1 treatment^47^. The mechanisms of these effects remain to be seen, and may relate to the expression of immunomodulatory factors by the tumor, or the re-shaping of the metabolic microenvironment, for example. To elucidate such pathways, close study of the tumor on a broader molecular scale, in conjunction with the immune cell composition and state, is necessary.

A further, surprising result from our bulk RNA analyses was that the LCAM axis was highly consistent with a change in fibroblast phenotype based on signatures derived from scRNAseq of NSCLC stromal clusters^5^. This association could suggest that the development of the tumor fibroblast phenotype is in response to overwhelming immune activation that may be instigated by an adaptive, antigen-specific response.

An important limitation of these findings relates to the site of initiation of the LCAM response; while the LCAM cellular module consists of cells undergoing an apparent active immune response, this study does not demonstrate the extent to which the module is instigated or perpetuated *in situ* at the tumor lesion. Specifically, despite evidence of clonally expanding T_activated_ cells, it remains unclear whether these lineages are primed *in situ* versus in tumor-draining lymph nodes (TdLN). While understanding the timescale of the tumor specific response will always be challenging due to variation in patient presentation timelines, it will nevertheless be important to correlate the cell types and states present in the TdLN in order to determine whether the LCAM response depends on lymph node priming, as well as to develop a deeper understanding of the spatial dynamics of the LCAM cell types.

Overall, the model presented here identifies an immune activation signature, derived from definitions of immune phenotypes defined by single-cell RNA and CITEseq, as an integrator of tumor-associated antigen load and driver mutation status that is not related to overall immune content. We believe that this axis, therefore, can serve as a more direct measure of antigen-specific, anti-tumor immune activation compared to previously suggested immune readouts.

## METHODS

### Human subjects

Samples of tumor and non-involved lung were obtained from surgical specimens of patients undergoing resection at Mount Sinai Hospital (New York, NY) after obtaining informed consent in accordance with a protocol reviewed and approved by the Institutional Review Board at the Icahn School of Medicine at Mount Sinai (IRB Human Subjects Electronic Research Applications 10-00472 and 10-00135) and in collaboration with the Biorepository and Department of Pathology.

### Tissue processing

Tissues were rinsed in PBS, minced and incubated for 40 minutes at 37°C in Collagenase IV 0.25mg/ml, Collagenase D 200U/ml and DNAse I 0.1mg.ml (all Sigma). Cell suspensions were then aspirated through a 18G needle ten times and strained through a 70-micron mesh prior to RBC lysis. Cell suspensions were enriched for CD45^+^ cells by either bead positive selection (Miltenyi) per kit instructions or FACS sorting on a BD FACSAria flow sorter (as indicated in Table S1) prior to processing for scRNAseq or CITEseq.

### ScRNA- and TCR-seq

For each sample, 10,000 cells were loaded onto a 10X Chromium single-cell encapsulation chip according to manufacturer instructions. Kit versions for each sample are indicated in Table S1. Libraries were prepared according to manufacturer instructions. QC of cDNA and final libraries was performed using CyberGreen qPCR library quantification assay. Sequencing was performed on Illumina sequencers to a depth of at least 80 million reads per library.

TCRseq was performed using the Chromium Single Cell 5’ VDJ kit, following manufacturer’s instructions. For patients 695 and 706, cells were subject to a CD2+ bead enrichment (Miltenyi) instead of CD45+ enrichment prior to encapsulation.

### CITEseq

For each sample, cell suspensions were split and barcoded using “hashing antibodies”^48^ staining beta-2-microglobulin and CD298 and conjugated to “hash-tag” oligonucleotides (HTOs). Hashed samples were pooled and stained with CITEseq antibodies that had been purchased either from the Biolegend TOTALseq catalog or conjugated using the Thunder-Link PLUS Oligo Conjugation kit (Expedeon). Sample hashing schemes and CITEseq panels are detailed in Tables S1 and S2, respectively. Stained cells were then encapsulated for single-cell reverse transcription using the 10X Chromium platform and libraries were prepared as previously described^15^ with minor modifications. Briefly, cDNA amplification was performed in the presence of 2pM of an antibody-oligo specific primer to increase yield of antibody derived tags (ADTs). The amplified cDNA was then separated by SPRI size selection into cDNA fractions containing mRNA derived cDNA (>300bp) and ADT-derived cDNAs (<180bp), which were further purified by additional rounds of SPRI selection. Independent sequencing libraries were generated from the mRNA and ADT cDNA fractions, which were quantified, pooled and sequenced together on an Illumina Nextseq to a depth of at least 80 million reads per gene expression library and 20 million reads per ADT library.

### MICSSS

FFPE tissues were stained using multiplexed immunohistochemical consecutive staining on a single slide as previously described^22^. Briefly, slides were baked at 37°C overnight, deparaffinized in xylene, and rehydrated in decreasing concentrations of ethanol. Tissue sections were incubated in citrate buffer (pH6 or 9) for antigen retrieval at 95°C for 30 minutes, followed by incubation in 3% hydrogen peroxide and in serum-free protein block solution (Dako, X0909) before adding primary antibody for 1 hour at room temperature. After signal amplification using secondary antibody conjugated to streptavidin-horseradish peroxidase and chromogenic revelation using 3-amino-9-ethylcarbazole (AEC), slides were counterstained with hematoxylin, mounted with a glycerol-based mounting medium and scanned for digital imaging (Pannoramic 250 Flash III whole-slide scanner, 3DHISTECH). Then the same slides were successively bleached and re-stained as previously described^22^. Primary antibodies were: anti-human CD10 (200103, R&D systems), DC-Lamp (1010E1.01, Novus biologicals), pan-cytokeratin (AE1/AE3, Dako), PDPN (D@-40, Ventana), CD163 (10D6, Novus Biologicals) and PD-L1 (E1L3N, Cell Signaling Tech).

### Analysis of Sequencing data

Transcriptomic and TCR library reads were aligned to the GRCh38/84 reference genome and quantified using Cellranger (v3.1.0). CITEseq ADT and CITEseq HTO reads were queried for antibody- and cell-specific oligonucleotide sequence barcodes in the designated read positions, including antibody sequences within a Hamming distance of 1 from the reference, using the feature-indexing function of Cellranger. Resulting alignment statistics are reported in Table S3. TCR data was aligned using Cellranger *vdj* function with default parameters.

### CITEseq processing and normalization

Doublets were removed based on co-staining of distinct sample-barcoding (“hashing”) antibodies ([*maximum HTO counts*]/[*2*^*nd*^ *most HTO counts*] < 5) and cell barcodes with few HTO counts (*maximum HTO counts* < 10) were also excluded. Cells were then assigned to samples based on their maximum staining HTO. HTO to sample associations are detailed in Table S1.

To normalize ADT counts across experimental batches given different CITEseq staining panels and sequencing runs, we performed a quantile-normalization on the ADT count values for each surface marker for the immune cells in each 10X encapsulation batch. To do this, the geometric average of the quantile function was computed across batches

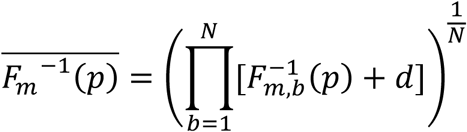

where 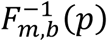 is the quantile function, or inverse cumulative distribution function, for counts of CITEseq marker *m* on immune cells in each of *N* 10X encapsulation batches *b* and regularization factor *d*=1 ADT count, evaluated at quantile *p* in interval [0,1]. This geometric average quantile function provided a reference function for a common mapping of cells based on their single-channel, batch-specific staining quantile *p* to a normalized staining intensity. Of note, this normalization method preserved the relationships between channels while constraining the observed differences in staining across experiments within individual channels.

### Unsupervised batch-aware clustering analysis

Immune cells from tumor and nLung samples were filtered for cell barcodes recording > 500 UMI, with < 25% mitochondrial gene expression, and with less than defined thresholds of expression for genes associated with red blood cells and with epithelial cells (Table S4). Cells were clustered using an unsupervised batch-aware clustering method we have recently described^16^ with minor adjustments. This EM-like algorithm, which was also based on earlier studies^49,50^, iteratively updates both cluster assignments and sample-wise noise estimates until it converges, using a multinomial mixture model capturing the transcriptional profiles of the different cell-states and sample specific fractions of background noise. We clustered 19 nLung and 22 tumor samples jointly and 46 additional tumor and nLung samples were mapped onto the final model as described

below.

The model definitions and estimation of model parameters were as described in (^16^). Specifically, the probability of observing gene *i* in cell *j* is defined as:

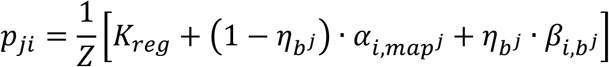

Where *map*^*j*^ and *b*^*j*^ are assignments of cells *j* to cell-type and batch respectively; 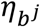 is the fraction of UMIs contributed by background noise; 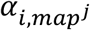 is the probability that a molecule drawn from celltype *map*^*j*^is of gene i (assuming no background noise) 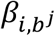 is the probability that a noise UMI drawn from batch *b*^*j*^ will be of gene *i*, and *K*_*reg*_ is a small regularization constant.

We also used here the pseudo expectation-maximization (EM) algorithm^16^ to infer the model parameters with minor modifications: (1) training set size was 2000 instead of 1000 cells and (2) the best clustering initiation was selected from 1000 instead of 10000 kmeans+ runs. For this clustering we included barcodes with more than 800 UMIs and used *K*_*reg*_*ds*_ = 0.2 ; (P_1,_P_2_) = (0^th^,30^th^) percentiles; *K*_*reg*_ = 5 · 10^−6^ ; k=60. Genes with high variability between patients were not used in the clustering. Those genes consisted of mitochondrial, stress, metallothionein genes, immunoglobulin variable chain genes, HLA class I and II genes and 3 specific genes with variable/noisy expression: *MALAT1, JCHAIN and XIST* (Table S4). Ribosomal genes were excluded only from the k-means clustering (Step 2.D as described in (^16^)). Samples used to generate this model included only those that were enriched for CD45+ immune cells using bead enrichment and were processed with the 10X Chromium V2 workflow.

### Integration of additional single-cell data

The resulting clustering model was used to analyze additional data that was both generated in-house or downloaded from public datasets. Single cells were mapped to clusters defined by the previously generated model *α*. Similarly to the clustering iterations, this process associates single-cells of a sample with multinomial probability vectors defined by the model and estimates the noise fractions of the sample *η*_*b*_ by optimizing the likelihood function (^16^):

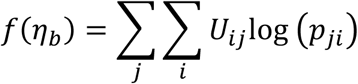

For *p*_*ji*_ as defined above, while 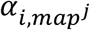 are updated using maximum likelihood.

Integrating inDrop data from (^6^) and 10X Chromium 5’ data required addressing the systematic differences^51^ in gene capture present between these technologies and 10X Chromium 3’ data that was used to develop the clustering model. Analysis of the differences in gene expression between the technologies suggested that a multiplicative correction factor *C*_*i*_ per each gene *i* could adjust for the capture efficiency differences. The following process was used to estimate the correction parameters:

1. Map cells to the original cluster models, as above, assuming absent noise in order to prevent the estimated noise term from being driven by error due to batch differences instead of true noise.
2. Re-calculate models using the average expression of the mapped cells for each cluster to form “data-based models” *α*^*D*^.
3. Calculate a weight matrix *W*, that weights individual genes for each cluster. *W* is calculated by

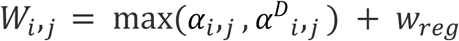

for original cluster model matrix *α*, data-based cluster model *α*^*D*^, gene *i*, cluster *j*, and regularization constant *w*_*reg*_ = 10^−10^. Since highly detected genes tend to dominate the mapping results, it is important to account for genes that are highly detected in either the original (10X Chromium V2) platform or the new platform
4. Construct a vector of gene-specific conversion factors that can operate between platforms:

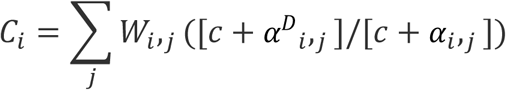

for regularization factor c = 10^−6^.
5. Generate transformed cluster models *α*’*i*,_*j*_ by multiplying the original models by the conversion vector and dividing by a normalization factor Z:

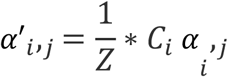
6. Map cells to transformed models without fixing the noise.

Analysis of the gene expression profiles of the mapped cells in each cluster demonstrated correspondence between the model and the mapped samples across the different technologies. (Fig S1B).

### Analysis of public datasets

Fastqs of scRNAseq data of tumors and nLung from 8 NSCLC patients^5^ acquired using 10X Chromium protocols was downloaded from ArrayExpress accessions E-MTAB-6149 and E-MTAB-6653. Sequencing reads were re-aligned using Cellranger as described above. Single-cells were mapped to clusters as described above. Tumor samples included 3 separate samples from the core, middle, and edge of each tumor. Regional tumor samples were considered separately for the intra-versus inter-patient variability analyses (Figure S1G, H). For remaining analyses, cell counts of projected tumor samples were pooled by patient.

scRNAseq data of tumors from 7 NSCLC patients^6^ acquired using inDrop was downloaded from GEO accession GSE127465. Since neutrophils were not detected in 10X Chromium data, cells that were annotated as neutrophils in the *GSE127465_human_cell_metadata_54773×25*.*tsv*.*gz* file were excluded from analysis. Cells were classified by projection as described above, using the modified procedure for inDrop data.

TCGA LUAD RNAseq data was downloaded using the *GDCquery* and *GDCdownload* functions from the *TCGAbiolinks* R package. *GDCquery* options included *project=“TCGA-LUAD”, data*.*category=“Transcriptome Profiling”, data*.*type=“Gene Expression Quantification”, workflow*.*type=“HTSeq – FPKM”, experimental*.*strategy=“RNA-Seq”*, and *legacy=F*. Whole exome sequencing data was downloaded using the *GDCquery_Maf* function with arguments *tumor=“TCGA-LUAD”* and *pipelines=“mutect2”*. Clinical data was downloaded using the *GDCquery_clinic* function with arguments *project=“TCGA-LUAD”* and *type=“clinical”*.

Processed CPTAC lung adenocarcinoma data was downloaded from the CPTAC Data Portal https://cptac-data-portal.georgetown.edu/cptacPublic/.

### Determination of sample-sample distances

Sample-sample distances were computed as the Euclidean distance between vectors consisting of the Log_10_-transformed cell type frequencies, where frequencies were computed as a fraction of total immune cells. A regularization factor of 10^−3^ was applied prior to applying the log-transform.

### Determination of myeloid cell type-specific gene scores

Lists of mutually-exclusive genes were used to compare monocytes, cDC2, and MΦ in Figure 2, and monocytes, AMΦ, and MoMΦ in Figure 3. For these analyses, genes were identified as “mutually exclusive” if the average expression was at least 2x greater in a given population than in the other comparison populations. To account for the large diversity of MoMΦ clusters, the maximum average expression of each MoMΦ subtype was used instead of the overall average expression. Resulting gene lists are presented in Table S4. Cells were scored according to the resulting gene lists as the Log-transformed fraction of UMI belonging to the gene list. Histograms were generated with the R function *density* using default parameters.

### Modules analyses

Gene-gene correlation modules were generated using a similar method to that previously described. Briefly, cells were downsampled to 2000 UMI prior to selecting a set of variable genes, similar to the selection of genes in preparation for seeding the clustering^16^. The gene-gene correlation matrix for this gene set was then computed for each sample over the cell population of interest. Correlation matrices were averaged following a Fisher Z-transformation. Applying the inverse transformation then resulted in the best-estimate correlation coefficients of gene-gene interactions across the dataset. Genes were clustered into modules using complete linkage hierarchical clustering over correlation distance. Histograms of module expression scores were generated with the R function *density* using default parameters.

### Classification of CD4+ versus CD8+ T_activated_ cells

CITEseq staining on a subset of patients was used to build a gene-set-based classifier that could use mRNA UMI data to discriminate CD4+ versus CD8+ cells within the T_activated_ cluster. To identify these gene sets, cells from 2 patients used as a training set were gated based on a Log_2_FC of raw ADT counts of raw CD4/CD8 > 1 and compared by differential expression. Genes were filtered by expression > 10^−4^ and a Log_2_FC > 1, and nonspecific or noise-related genes such as those associated with cell-cycle, long-non-coding RNAs, heat shock proteins, immunoglobulin genes, ribosomal proteins, *XIST*, and histone transcripts. Resulting gene lists are reported in Table S4. Cells were scored based on the fraction of RNAs belonging to the resulting gene lists, and a discrimination threshold for the ratio of the CD4 vs. CD8 gene lists was determined based on the overall accuracy in discriminating between CITEseq-defined CD4+ vs. CD8+ cells in the training set. This gene score discriminator was validated using cells from a test set comprised of cells from 4 additional patients analyzed by CITEseq (584, 593, 596, 630), and on cells with unique detection of either *CD4* or at least one of (*CD8A, CD8B*).

### Analysis of cycling T cell cluster

To analyze the phenotypic makeup of the cluster of T cells expressing cell-cycle genes, we generated gene sets based on the other T cell phenotypes described here to score each cell within the cluster. To do this, we pooled the cells of each other T cell phenotype to compute its average expression. We then identified a gene list for each phenotype defined by expression > 1e-5 and Log2FC > 0.25 compared to the maximum of the other phenotypes. From this list, we excluded variable TCR genes, and other genes associated with noise or cell stress. The gene lists for the T_naive/CM-like_ cell types were grouped, since these phenotypes were very similar. Resulting gene lists are reported in Table S4.

For each cell in the cycling cluster, we then computed the fraction of UMI belonging to each gene signature after removing UMIs belonging to the list of genes associated with the cycling cluster that was calculated as above. We performed spherical k-means clustering using the function *skmeans()* in the *skmeans* R package on these signature fractions in order to group cells within the cycling cluster according to phenotypic subtype by spherical k-means cluster.

### Single-cell TCRseq analysis

Single T cells were grouped by clonotype according to their precise combination of α and β chains present (uniquely defined by CDR3 sequence and V, D, and J gene usage), with the following acceptations in order to filter for high quality singlets:

1. Cells with contigs encoding > 3 productive α and β chains were excluded as multiplets.
2. Cells with contigs encoding > 3 productive α and β chains that completely overlapped with observed cells within the multiplets were also excluded as multiplets.
3. Remaining cells with 3 unique α and β chains that could be uniquely associated with similar cells displaying 2 unique α and β chains were assumed to be clonally related, whereas cells that could be similarly associated with multiple distinct sets of cells expressing 2 unique α and β chains were excluded as doublets.
4. Cells in which a single TCR chain was observed were assumed to be clonally related to any cells with 2 unique α and β chains to which they uniquely associated.
5. Remaining cells in which a single TCR chain was observed were excluded if they matched ambiguously to multiple cells with 2- or 3-chains.
6. Clonality scores were computed for each T cell type in each patient as *1-Peilou’s eveness* over the set of unique TCRs as previously described^52^.

### Ligand-receptor analysis

Ligand-receptor intensity scores for a set of secreted ligands (ref(^33^) and Table S6) were calculated as previously reported^16^. Briefly, for each ligand-receptor interaction, for each source cell type and each receiver cell type, the intensity score was equal to the product of ligand generation from the source cell type relative to the total RNA with the expression of the receptor on the receiver cell type. Scores were independently calculated for LCAM^hi^ and LCAM^lo^ patient sets in nLung and Tumor tissues. To determine these patient sets, patients were sorted by the geometric mean of lineage-normalized cellular frequencies of LCAM^hi^ and LCAM^lo^ cell types, and the top half of patients were defined as LCAM^hi^ with the bottom half defined as LCAM^lo^. Only patients analyzed using 10X Chromium V2 with immune cells purified with magnetic beads were used for this analysis. The patients included in these groups were: LCAM^hi^: (408, 403, 522, 371, 570, 714, 584, 377, 406, 564, 630, 578, 514); LCAM^lo^: (571, 596, 393, 593, 626, 378, 370, 410, 572, 558, 581, 596, 729).

### Identification of LCAM^hi^ and LCAM^lo^ bulk-RNA gene signatures

To define genes that could probe the presence of LCAM^hi^ or LCAM^lo^ cell types in bulk RNA data, we adopted a similar strategy to that used previously for the projection of bulk data onto signatures defined by cellular axes as measured with scRNA^16,34^. Cells were evenly sampled from LCAM^hi^ and LCAM^lo^ patients (1409 cells per patient), and sampled cells were then pooled within the groups. Differentially expressed genes (*FDR*<10^−3^ and Log2FC > 1) were retained. Genes that were expressed in the filtered epithelial cells > 2x higher than in immune cells on average were removed. Among the remaining differentially expressed genes, those that were expressed on average within any LCAM^hi^ or LCAM^lo^ subtype with Log2FC > 3 compared to the highest expressing subtype in the opposite group were retained. These gene lists were further abbreviated to include no more than 10 genes per cell type, in order to balance the number of genes coming from any individual cell type. In order to increase the differential expression effect sizes observed, only the most extreme 6 LCAM^hi^ and LCAM^lo^ patients processed with CD45+ bead enrichment and 10X Chromium V2 were included in the differential expression analysis. These patients were LCAM^hi^: (408, 403, 714, 522, 371, 570), and LCAM^lo^: (571, 596, 393, 593, 626, 378).

### Calculation of LCAM^hi^, LCAM^lo^, and ensemble LCAM scores in bulk-RNA datasets

Bulk RNA expression datasets were log-transformed and z-scored. For each cell type associated with LCAM^hi^ or LCAM^lo^, the resulting z-scores of the associated genes were averaged and z-scored. A summary average of these values was then computed across all the cell types associated with either LCAM^hi^ or LCAM^lo^ cell types.

### Published statistics for TCGA Lung adenocarcinoma patients

Estimates of total immune content present in each TCGA sample (ESTIMATE score)^35^ were download from https://bioinformatics.mdanderson.org/public-software/estimate/. Scores associating mutational signatures^41^ with individual TCGA samples were downloaded from the mSignatureDB^42^ website http://tardis.cgu.edu.tw/msignaturedb/. For the present study, detection of Signature 4 was used to indicate presence of smoking-related mutations.

Counts of Indel Neoantigens, SNV Neoantigens, and CTA score in TCGA cases were accessed from Table S1 of ref. (^53^).

### Generation of stromal cell type scores

Fibroblast and endothelial cell count matrices from tumor and nLung of 8 NSCLC patients^5^ were downloaded from https://gbiomed.kuleuven.be/english/research/50000622/laboratories/54213024/scRNAseq-NSCLC. Previously-applied^5^ cluster annotations were assumed, where endothelial cluster 6 was defined as “lymphatics”, and endothelial and fibroblast clusters were defined based on enrichment in tumor or nLung: endothelial clusters 3 and 4 were pooled as “Tumor BEC”, endothelial clusters 1 and 5 were pooled as “Normal BEC”, fibroblast cluster 1 was defined as “Normal fibroblast”, and fibroblast clusters 1, 2, 3, 4, 5, and 7 were pooled as “CAF”. For each of these cell types, gene scores were defined based on a minimum average expression of 10^−4^ and a minimum fold-change threshold of 4 compared to any other stromal cell type. Cell type gene-scores were defined in TCGA lung adenocarcinoma using the average z-scored gene expression of each stromal gene list.

## DATA AVAILABILITY

Human scRNAseq, TCRseq, and CITEseq data is available at GEO accession GSE154826.

## ACKNOWLEDGMENTS

This work was supported by National Institutes of Health (NIH) grants 5T32CA078207 (to A.M.L.). We thank A. Magen, P. Hamon, M. Casanova-Acebes for critical comments on the manuscript; and the Mount Sinai flow cytometry core, Human Immune Monitoring Center and Mount Sinai Biorepository for support. Research reported in this paper was supported by the Office of Research Infrastructure of the National Institutes of Health under award numbers S10OD018522 and S10OD026880. The content is solely the responsibility of the authors and does not necessarily represent the official views of the National Institutes of Health. Data used in this publication were generated by the TCGA Research Network, and the National Cancer Institute Clinical Proteomic Tumor Analysis Consortium (CPTAC). Research support was provided by Regeneron and Takeda. We recognize the patients and their families for their important contributions and sacrifices.

## AUTHOR CONTRIBUTIONS

MM and EK conceived the project. AML, AR, and MM designed the experiments. AML, EK, and MM wrote the manuscript. AML, EK, and MD performed computational analysis. TM, MB, AW, and RF facilitated access to human samples. AML, JG, CC, BM, AT, LW, JL, NM, GM, and KT performed experiments. NRD and GT funded part of the study. AL conducted patient consents and facilitated regulatory items. JG, JM, GM, ZZ, FP, RS, AK, PW, HS, and TM provided further intellectual input.

## FIGURE LEGENDS

**Figure S1.**
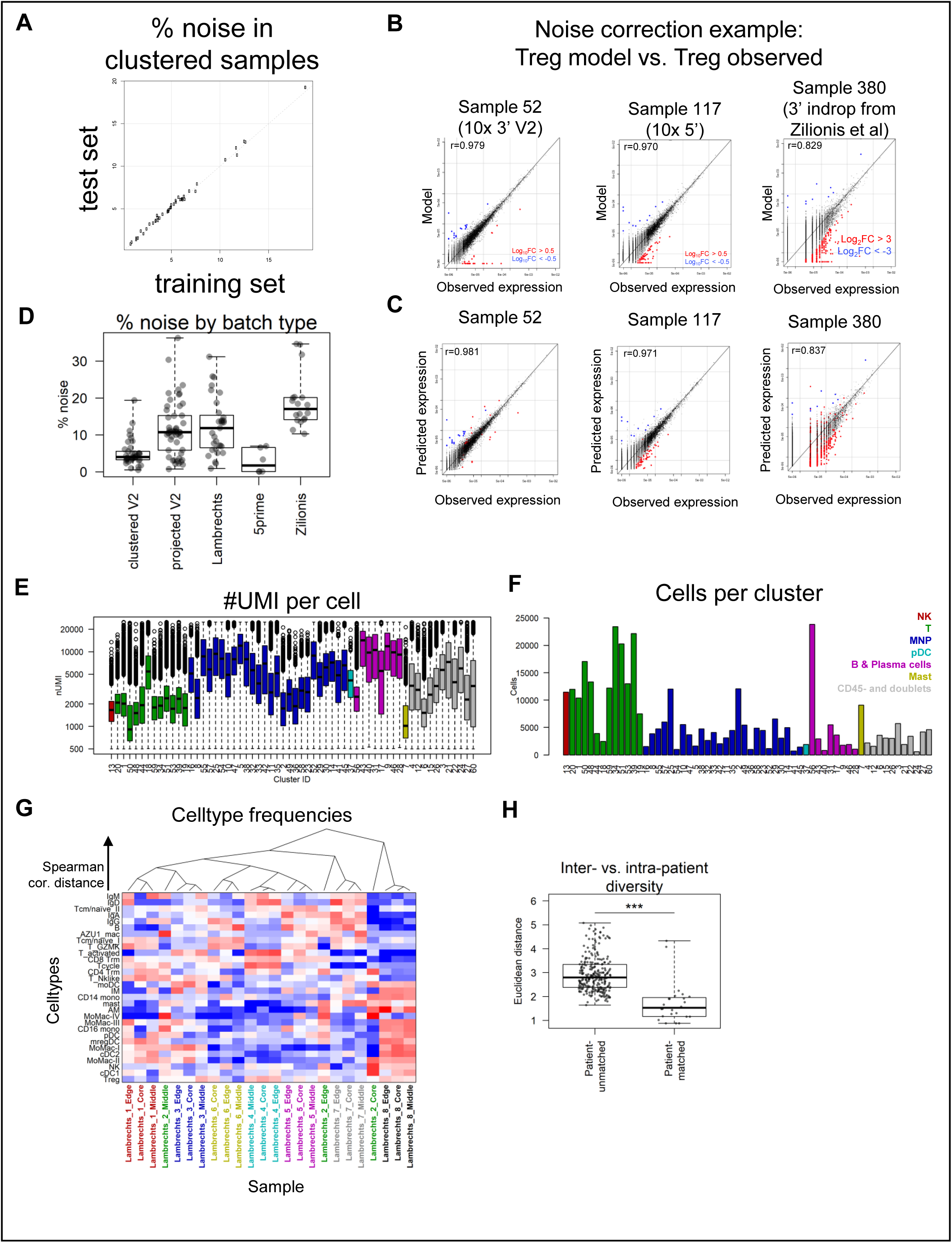
Integration of scRNA samples and datasets for common cell type analysis. **A**, Comparison of per-sample estimated noise levels in the training set of cells used for clustering and model formation (x-axis) compared to the per-sample estimated noise in a withheld test set of cells that were mapped to the model clusters by probabilistic projection. **B and C**, Illustration of how incorporating a fit noise component improves the concordance between predicted expression and of cells mapped to the T_reg_ cluster and observed expression. Y-axis shows the predicted expression of T_regs_ in individual samples without accounting for noise (**B**) or accounting for noise (**C**), against the observed average expression (X-axis). Genes were color-coded by the ratio between the observed expression and the model without accounting for noise. Estimation of the noise component is detailed in the methods. **D**, Per-sample estimated noise levels in 10X chromium V2 samples that were used for clustering, 10X chromium V2 samples that were analyzed by projection onto the clustering model and not used in the clustering, 10X chromium 5’ samples that were analyzed by projection, and samples from external datasets^5,6^ that were analyzed by projection. **E**, Boxplots showing the distribution of UMI per cell in each cluster. **F**, Barplots showing number of cells in Mount Sinai dataset mapping to each cluster. **G**, Heatmap showing row-normalized cell type frequencies in a public dataset^5^ with samples spanning 3 regions each in a cohort of 8 NSCLC patients. Samples are clustered by spearman correlation distance. Sample names are colored by patient. **H**, Box plots of Euclidean distances based on log-transformed cluster frequencies between samples of different patients or from the same patient, as in (**G**), from ref. ^5^. *** P < 0.001, Wilcoxon rank-sum test.

**Figure S2.**
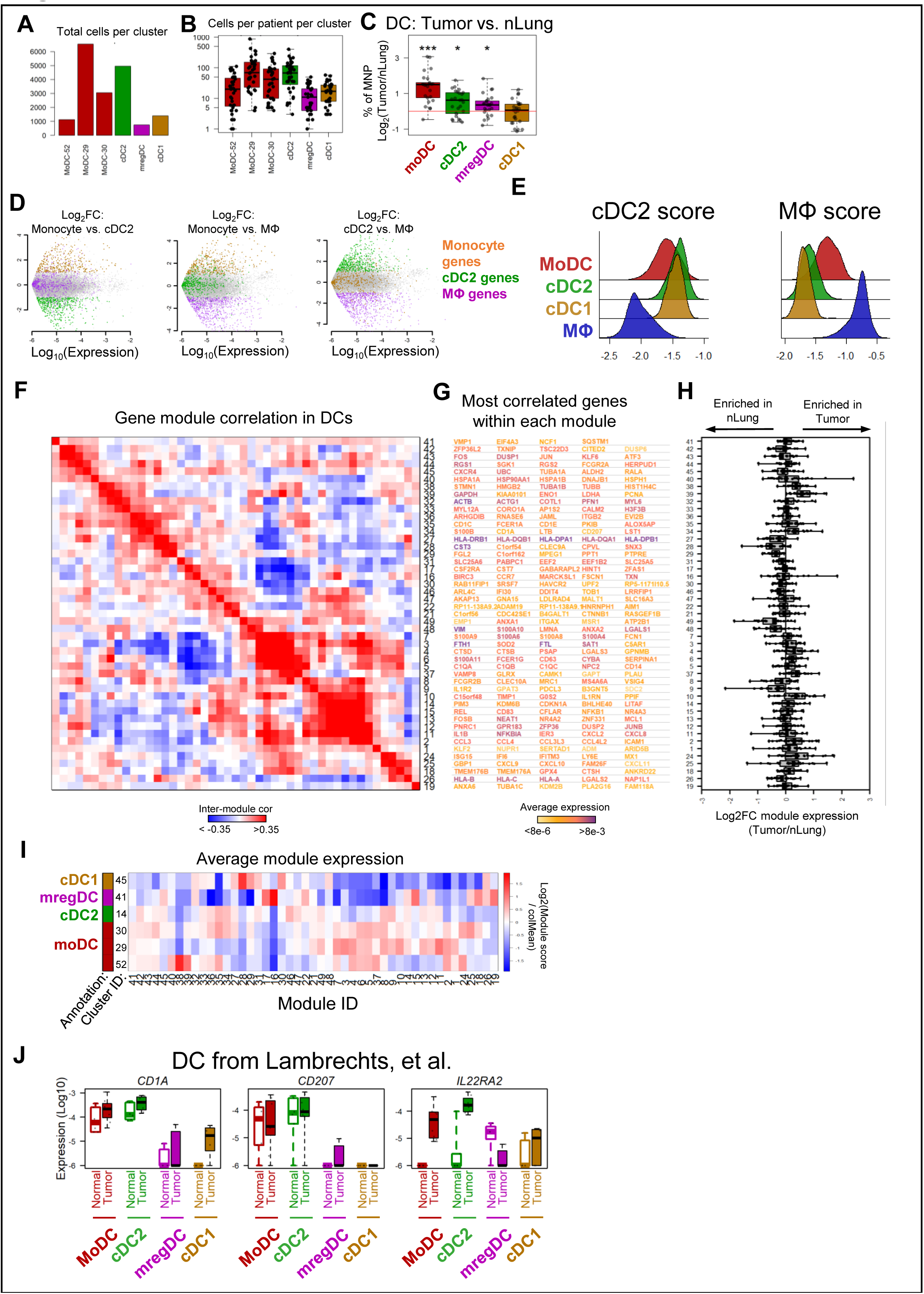
Module analysis of DC. **A**, Barplots showing total number of cells mapped to each individual DC cluster in the Mount Sinai cohort. **B**, Boxplots showing number of cells mapped to each individual DC cluster per tumor sample in the Mount Sinai cohort. **C**, Differences between tumor and nLung of DC frequencies normalized to total MNP; *P<0.05, **P<0.01, ***P,0.001 (Wilcoxon signed-rank test with Bonferroni correction; N=26 matched tissue pairs with >250 MNP observed in each tissue). **D**, Log_2_FC and expression level distributions of gene sets that are mutually exclusively expressed in CD14+ monocytes, MΦ, and cDC2 (See Figure 2G). **E**. Histograms of cDC2 and MΦ scores, using gene lists generated as shown in (**D**). **F-I**, Gene module analysis of DC clusters. Correlation of gene module expression across all DC (**F**), five example genes from each module, ranked by correlation to the other genes in the module and colored by total expression in DC (**G**), boxplots showing Log_2_FC of module expression among all DC between patient matched tumor and nLung samples (**H**), and normalized average cluster expression of modules (**I**). **J**, Boxplots showing expression of LCH-like signature genes across DC populations in distinct nLung and tumor samples from ref. ^5^.

**Figure S3.**
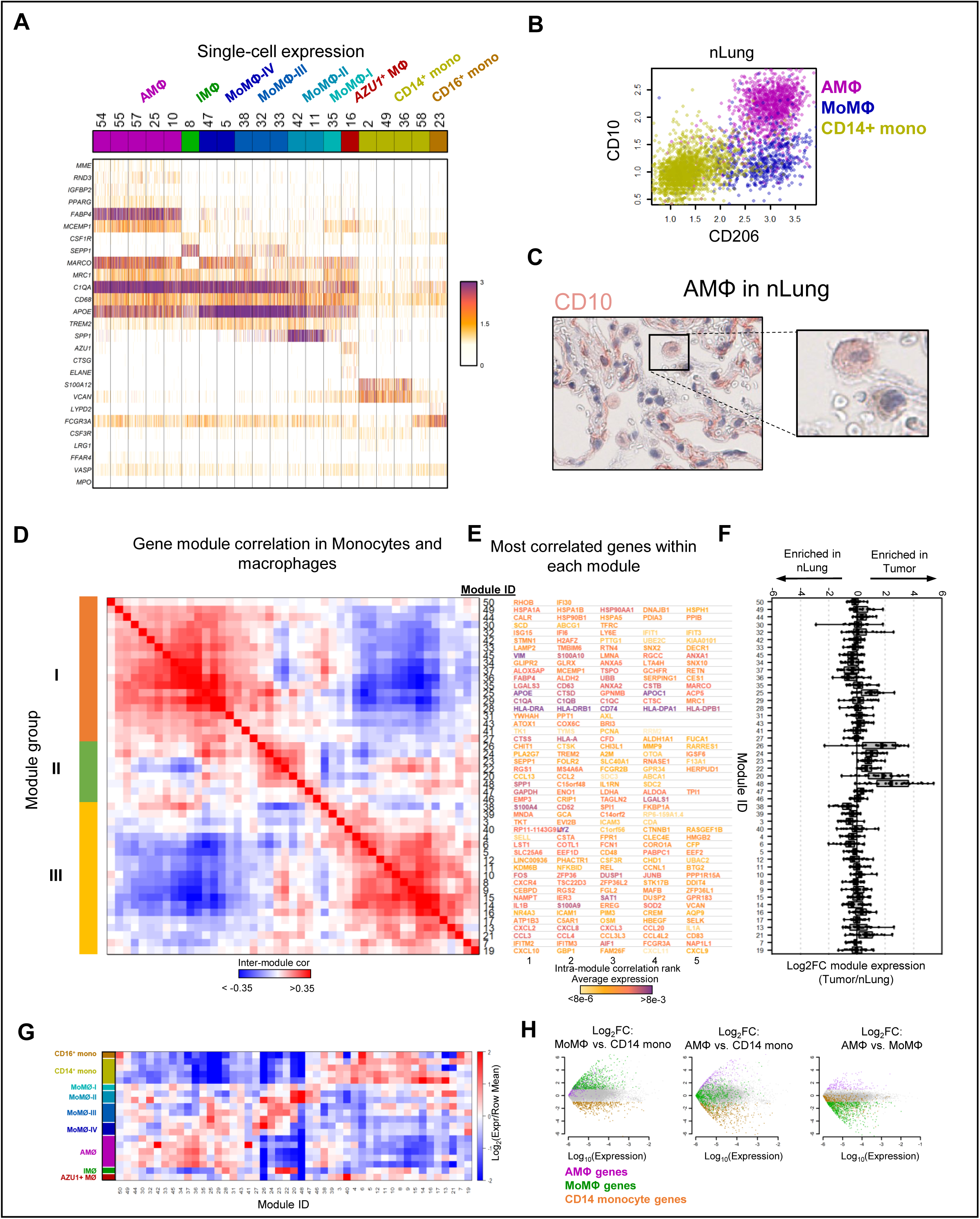
Diversity of nlung and tumor-infiltrating MΦ populations. **A**, Expression of key genes discriminating scRNAseq clusters of monocytes and MΦ, grouped by cell type annotation. Heatmap shows the number of UMI per cell. Clusters are shown using an even number of randomly selected cells from each, drawing from 35 tumor and 32 nLung samples. Cells were downsampled to 2000 UMI/cell. **B**, Scatter plots showing normalized CITEseq CD10 and CD206 surface marker counts on AMΦ, MoMΦ, and CD14+ monocytes in nLung of a representative patient. **C**, IHC of CD10 staining AMΦ in the airspaces of nLung tissue. **D-G**, Gene module analysis of monocyte and MΦ clusters. Correlation of gene module expression across all monocytes and MΦ (**D**). Module groups illustrate groups of correlated modules which are expressed most specifically on AMΦ, MoMΦ, and monocytes (see **G**). Five example genes from each module, ranked by correlation to the other genes in the module and colored by total expression in monocytes and MΦ (**E**), boxplots showing Log_2_FC of module expression among all monocytes and MΦ between patient matched tumor and nLung samples (**F**), and normalized average cluster expression of modules (**G**). **H**, Log_2_FC and expression level of gene sets that are mutually exclusively expressed in CD14+ monocytes, AMΦ, and MoMΦ (See Figure 3D-F).

**Figure S4.**
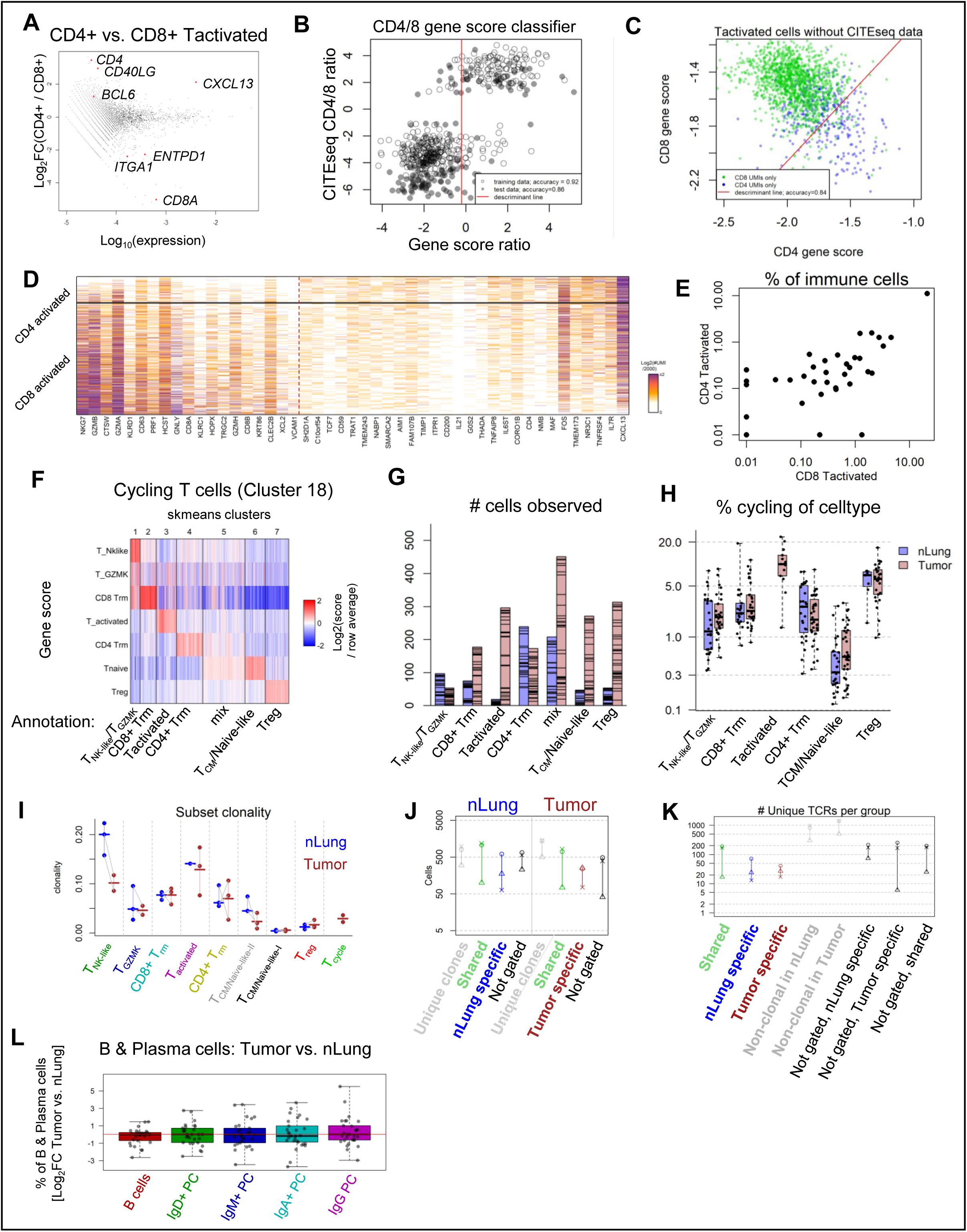
Phenotypic dissection of activated, cycling, and clonally expanded T cells. **A**, Differential expression within the T_activated_ cluster of cells staining for CD4 versus CD8 by CITEseq (y-axis) vs. average T_activated_ expression (x-axis). **B**, Classification of T_activated_ cells as CD4+ or CD8+ based on the ratio of CD4-related or CD8-related gene signatures learned from cells of 2 patients (training set; open circles) and validated on cells of 4 additional validation patients (test set; black dots). Red line indicates gene ratio threshold learned from the training set. Only cells where the CITEseq CD4:CD8 count ratio is >2 or <1/2 are considered. **C**, Validation of CD4/CD8 classification scheme shown in (**B**) for cells without CITEseq staining. Cells were considered to be CD4+ or CD8+ based on unique RNA detection of either *CD4* (blue points) or at least one *CD8A* or *CD8B* transcript (green points). The discriminant line is equivalent to the gene ratio threshold learned from CITEseq data, shown in (**B**). **D**, Expression of key genes in CD4-related and CD8-related gene signatures for discriminating CD4+ and CD8+ activated T cells. Cells are sorted by ratio of these gene signatures, and the line is drawn to indicate the cells discriminated based on the threshold in panel (**B**). Heatmap shows the number of UMI per cell. Cells represent T_activated_ cells from 35 tumors, and were downsampled to 2000 UMI/cell. **E**, Frequency of CD8+ or CD4+ T_activated_ cells across 35 patients, as determined by gene signature scores learned from CITEseq (as in **A-D**). **F-H**, Spherical k-means sub-clustering on cell type scores of cells within the cycling T cell cluster 18 based on gene scores generated from other T cell clusters. Heatmap of single-cell expression of cell type scores, grouped by sub-cluster (**F**), number of cells in each sub-cluster (**G**; nLung shown in blue, tumor in brown; lines dividing bars horizontally discriminate groups of cells from distinct patients), and the frequency of cycling T cells of each T cell phenotype (**H**; data points represent samples with at least 50 cells of the given phenotype). **I**, TCR clonality score of phenotypic groups in nLung (blue) and tumor (brown). Dots represent individual samples with at least 30 cells of indicated phenotype. N=3 patients with tumor-nLung pairs. **J**, Number of cells within each TCR category, determined as in Figure 5D, in matched nLung and tumor samples of 3 patients, each patient indicated by shape. **K**, Number of unique TCRs represented in each TCR category, determined as in Figure 5D. **L**, Differences between tumor and nLung of lineage-normalized B and plasma cell type frequencies. All comparisons were not significant (P>0.05, Wilcoxon signed-rank test, N=32 matched tissue pairs).

**Figure S5.**
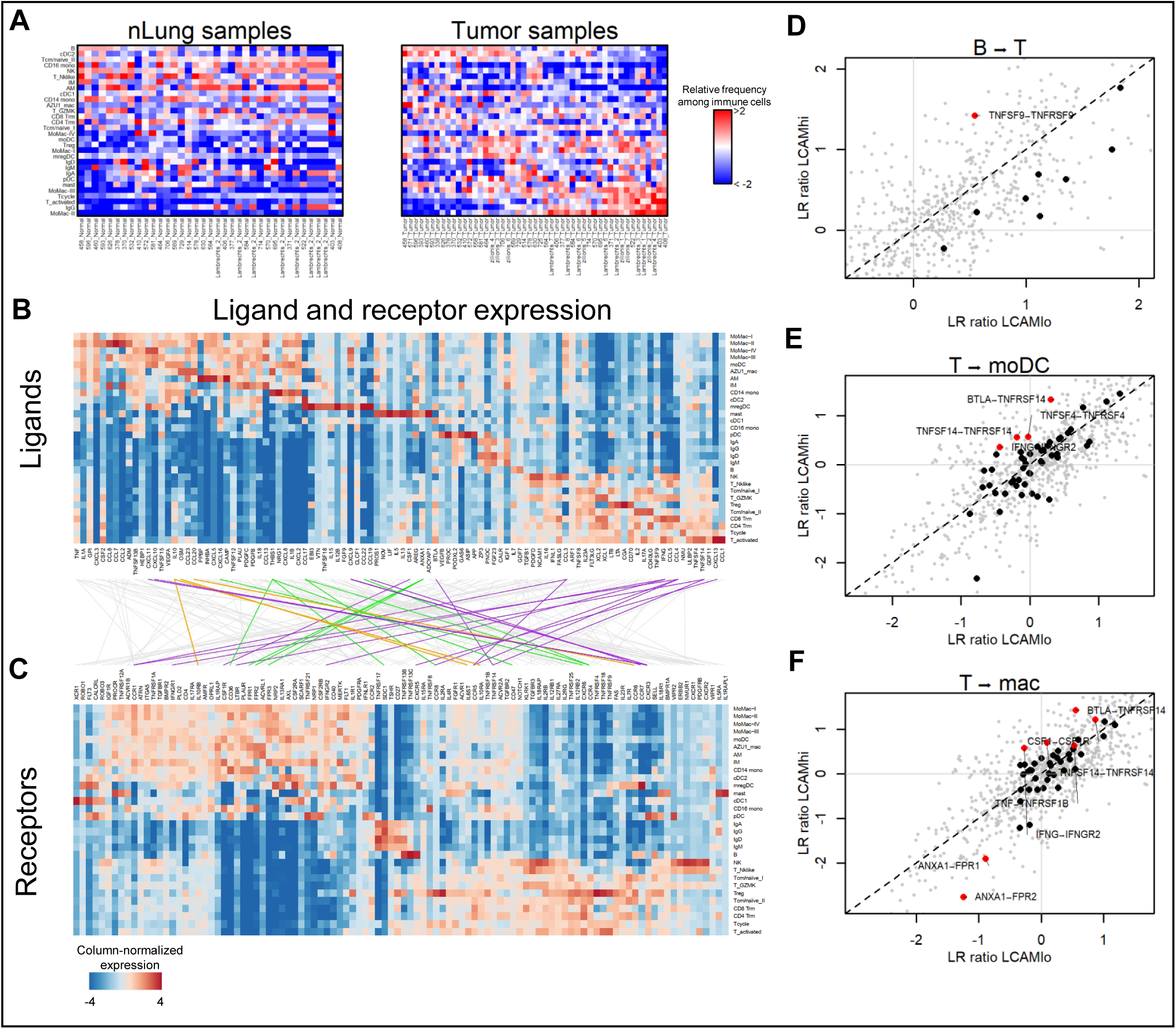
Ligand-receptor intaractions in LCAM^hi^ and LCAM^lo^ tumors. **A**, Lineage-normalized cell type frequencies of all cell types among pooled nLung and Tumor samples from Mount Sinai and refs. ^5,6^ (50 tumor patients with 40 matched nLung samples). **B and C**, Column-normalized expression of highly expressed secreted ligands (**B**) and associated receptors (**C**) across all immune cell types, connected by lines linking ligands to receptors. Connectors are colored by association with LCAM^hi^ patients (purple), LCAM^lo^ patients (green), or all tumors (orange). **D-F**, Log2 Ratio of ligand-receptor (LR) intensity scores between tumor and nLung of LCAM^hi^ patients, (“LR ratio”; y-axis) and LCAM^lo^ patients (x-axis) as in Figure 5D, highlighting in bold LR ratios for interactions between B cell ligands and T cell receptors (**D**), T cell ligands and MoDC receptors (**E**), and T cell ligands and MΦ receptors (**F**). Labelled interactions are plotted in red.

**Figure S6.**
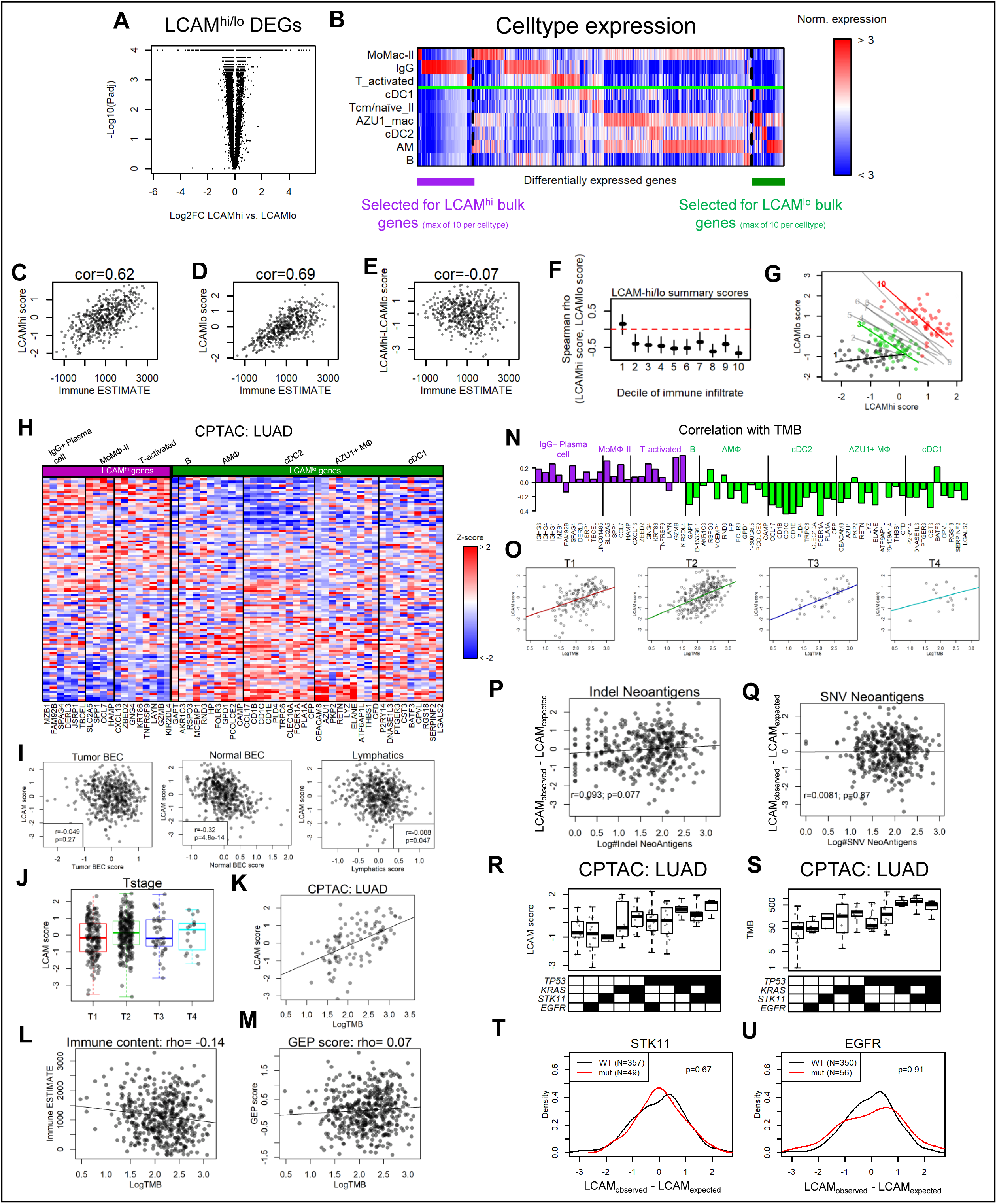
Projection of bulk RNA samples onto signatures defined by the LCAM scRNA axis. **A and B**, Derivation of the LCAM^hi^ and LCAM^lo^ gene signatures for scoring bulk RNA samples. Identification of differentially-expressed genes between averaged scRNAseq samples of LCAM^hi^ and LCAM^lo^ patients (**A**), and identification of differentially expressed genes that are specific to genes in the LCAM^hi^ or LCAM^lo^ cell types (**B**). **C-E**, Scatter plots of immune ESTIMATE score^35^ with the LCAM^hi^ signature score (**C**), the LCAM^lo^ signature score (**D**), or the difference between the LCAM^hi^ and LCAM^lo^ signature scores (i.e. the ensemble LCAM score; **E**). **F**, Spearman correlation of the LCAM^hi^ and LCAM^lo^ signature scores among the deciles of immune content. Error bars represent the 95% confidence interval around the estimate of the spearman correlation. **G**, Scatter plots of the LCAM^hi^ and LCAM^lo^ signature scores, showing the 1^st^ (black), 3^rd^ (green), and 10^th^ (red) deciles of immune content. Labelled trend lines are shown for other deciles. **H**, Normalized expression of LCAM^hi^ and LCAM^lo^ bulk-RNA signature genes in the CPTAC lung adenocarcinoma dataset. **I**, Scatter plots of the ensemble LCAM score (y-axis) with signature scores based on genes that are specific for tumor blood endothelial cells (BEC; left), normal BEC (center), and lymphatic endothelial cells. Stromal signatures are based on the stromal data reported in ref. ^5^. **J**, Boxplots showing ensemble LCAM scores among TCGA lung adenocarcinoma patients by TNM T-stage. **K**, Scatter plot of LogTMB and ensemble LCAM score in CTPAC lung adenocarcinoma patients, with linear regression line. **L and M**, Scatter plots of the LogTMB and immune ESTIMATE score^35^ (**L**) and the T-cell inflamed gene expression profile (GEP^39,40^; **M**) in TCGA lung adenocarcinoma patients. **N**, Correlation between individual genes comprising the LCAM^hi^ and LCAM^lo^ bulk gene signatures and LogTMB in TCGA lung adenocarcinoma patients. **O**, Scatter plots of LogTMB and the LCAM ensemble score for patients by T-stage. **P and Q**, Scatter plots of the number of indel-induced neoantigens (**P**) and SNV-induced neoantigens (**Q**) as computed in ref. ^53^, and the residuals of the regression of the ensemble LCAM score on the LogTMB. **R and S**, Boxplots showing either the ensemble LCAM score **G**), or TMB (**H**) among CPTAC lung adenocarcinoma patients, divided by combinations of mutated driver mutations. **T and U**, Histograms of residuals of the regression of the ensemble LCAM score on the LogTMB, with patients stratified by *STK11* (**T**) or *EGFR* (**U**) mutational status (Two-sided t-test).

## SUPPLEMENTAL TABLES

**Table S1**. Sample table, with information about patient, tissue, 10X loading, and QC metrics

**Table S2**. CITEseq panels used

**Table S3**. QC table of GEX, HTO, and ADT libraries

**Table S4**. Gene lists used in paper

**Table S5**. Gene modules

**Table S6**. Ligand-receptor pairs used in the analysis

**Table S7**. Ligand-receptor statistics

